# Reversible restriction of electron flow across cytochrome *b_6_f* in dark acclimated cells limited for downstream electron sinks

**DOI:** 10.1101/2022.10.04.507358

**Authors:** Shai Saroussi, Devin Karns, Dylan C. Thomas, Petra Redekop, Tyler M. Wittkopp, Matthew C. Posewitz, Arthur R. Grossman

**Affiliations:** The Carnegie Institution for Science, Department of Plant Biology, Stanford, CA 94305, USA; Department of Chemistry and Geochemistry, Colorado School of Mines, Golden, CO 80401, USA; Stanford University, Department of Biology, Stanford, CA 94305, USA

**Author notes:** Corresponding authors: Arthur Grossman, Address: Department of Plant Biology, Carnegie Institution for Science, 260 Panama St., Stanford, CA 94305, USA, And, Shai Saroussi, Address: Department of Plant Biology, Carnegie Institution for Science, 260 Panama St., Stanford, CA 94305, USA.

## Abstract

Photosynthetic organisms frequently experience abiotic stresses that restrict their growth and development. Under such circumstances, most absorbed solar energy cannot be used for CO_2_ fixation and can cause the photoproduction of reactive oxygen species (ROS) that can damage the photosynthetic reaction centers, photosystems I and II (PSI and PSII), resulting in a decline in primary productivity. This work describes a biological ‘switch’ in the green alga *Chlamydomonas reinhardtii* that reversibly restricts photosynthetic electron transport (PET) at the cytochrome *b*_*6*_*f* complex when reductant and ATP generated by PET are in excess of the capacity of carbon metabolism to utilize these products; we specifically show a restriction at this switch when *sta6* mutant cells, which cannot synthesize starch, are limited for nitrogen (growth inhibition) and subjected to a dark-to-light transition. This restriction, which may be a form of photosynthetic control, causes diminished electron flow to PSI, which prevents PSI photodamage. When electron flow is blocked the plastid alternative oxidase (PTOX) may also become activated, functioning as an electron valve that dissipates some of the excitation energy absorbed by PSII thereby lessening PSII photoinhibition. Furthermore, illumination of the cells following the dark acclimation gradually diminishes the restriction at cytochrome *b*_*6*_*f* complex. Elucidating this photoprotective mechanism and its modulating factors may offer new insights into mechanisms associated with photosynthetic control and offer new directions for optimizing photosynthesis.

## INTRODUCTION

Oxygenic photosynthesis is a fundamental biological process performed by plants, algae, and photosynthetic bacteria that supports most life on Earth. The photosynthetic complexes absorb light, which provides the energy to oxidize water, using the extracted electrons to generate NADPH, reduced ferredoxin (FDX), and energy as ATP and electrochemical gradients. The ATP and NADPH fuel downstream CO_2_ fixation via the Calvin-Benson-Bassham cycle (CBBC) (**Figure 1A**), which produces the carbon (C) backbones required for nutrient assimilation, the generation of essential cellular building blocks, respiratory substrates, and C polymers (e.g., starch and lipids) that serve in energy storage. Coordination between PET and C metabolism involves ‘redox control’ of CBBC enzymes, which is primarily mediated by FDX and thioredoxin (TRX) (Buchanan, 2016). In the light, as the redox potential in the chloroplast becomes more negative, TRX reduces disulfide bonds on targeted CBBC enzymes leading to an increase in their activity by up to 40-fold, while oxidation of these disulfides when the redox potential becomes more positive results in diminished enzyme activity (Michelet et al., 2013; Wirtz et al., 1982) (light blue box, upper right in **Figure 1A**).

**Figure 1:**
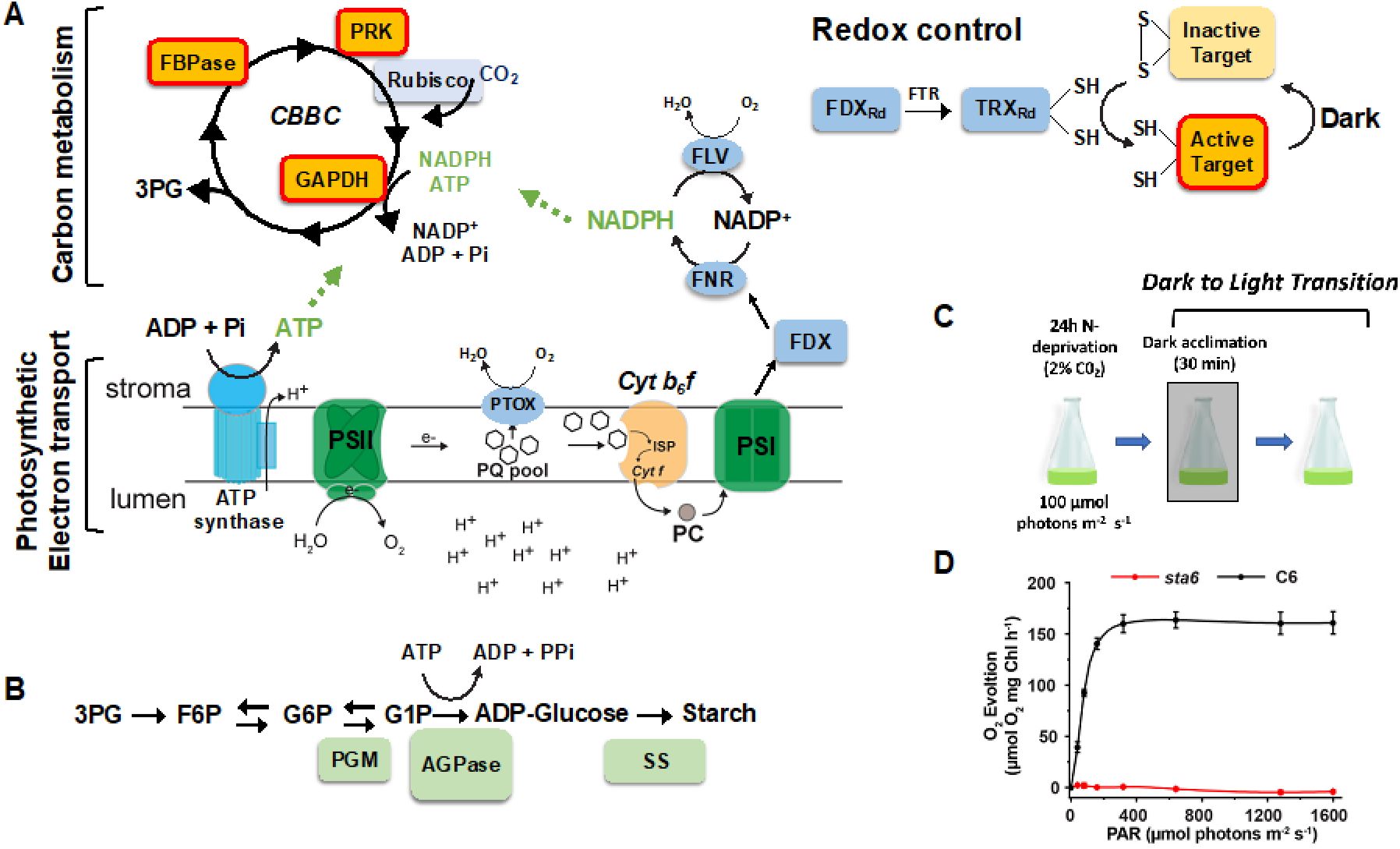
Key aspects of the photosynthetic system and experimental design: **A**. Schematic representation of the link between PET and downstream C metabolism (CBBC; upper left). Electrons extracted from H_2_O by PSII flow through the electron transport chain generating a proton (H^+^) gradient across the thylakoid membranes and ultimately reducing NADP^+^ to NADPH. The energy of the H^+^ gradient is used to synthesize ATP by the ATP synthase. CBBC utilizes both ATP and NADPH (green lettering and dashed green arrows) to fix CO_2_ by Rubisco (light blue box). The CBBC includes several enzymes that are activated by redox control (activated enzymes highlighted in orange boxes with red frames). Upper Right: Redox activation of targeted CBBC enzymes by reduction of specific disulfide bonds, which is mediated by FDX and TRX systems. Activation is reversible upon oxidation of the disulfide bonds in the dark. **B**. Starch biosynthesis: Metabolites of the starch synthesis pathway are in black lettering. Starch metabolism-related enzymes are highlighted by a light green rectangular background. **C**. Experimental design: After growing the cells mixotrophically in moderate light (100 µmol photons m^−2^s^−1^) to mid log phase, they were transferred to photoautotrophic medium devoid of N and acclimated for 24 h in moderate light. Following a 30 min dark acclimation period, the cells were exposed to light (intensities indicated in text). **D**. Net O_2_ evolution of N-deprived C6 (black curve) and *sta6* (red curve) cultures. N=3 ± SE. Abbreviations: PSI - photosystem I; PSII - photosystem II; *Cyt b*_*6*_*f* - cytochrome *b*_*6*_*f*; ISP - iron sulfur protein; *Cyt f* - cytochrome *f*; PC - plastocyanin; PTOX - plastid terminal oxidase; FLV, flavodiiron protein; FDX - ferredoxin; FNR - ferredoxin-NADPH oxidoreductase; FTR - ferredoxin-thioredoxin reductase; Rubisco - ribulose-1-5-bisphosphate carboxylase-oxygenase; GAPDH - glyceraldehyde 3-phosphate dehydrogenase; PGM - phosphoglucomutase; AGPase - ADP-glucose pyrophosphorylase; SS - starch synthase; FBPase - fructose-1,6-bisphosphatase; PRK - phosphoribulokinase. Metabolites: 3PG - 3-phospho-glyceraldehyde; F6P - fructose-6-phosphate; G6P - glucose-6-phosphate; G1P- glucose-1-phosphate. PAR - photosynthetically active radiation.

In nature, environmental change and abiotic stress can cause an imbalance in the activities of PET and the CBBC. Under conditions in which the photosynthetic machinery is unable to effectively balance photosynthetic electron output with utilization, light energy may be converted to toxic ROS that can damage photosystems I (PSI) and II (PSII) through a process known as photoinhibition, and modify other cellular components including DNA and lipids, potentially leading to cell death. Mechanisms have evolved to protect the cell from photodamage through the dissipation of excess absorbed light energy as heat in a process known as nonphotochemical quenching (NPQ) (Müller et al., 2001; Rochaix, 2014). NPQ and other mechanisms that reduce the damaging impact of excess absorbed light energy are highly relevant to both ecological and agricultural issues as anthropogenic activities are escalating the rate of climate/environmental change (Malnoë, 2018; Müller et al., 2001), which can markedly impact photosynthetic function. An excess of energetic photosynthetic electrons, however, can also be dissipated by photochemical outlets that involve the reduction of oxygen (O_2_) to regenerate water (H_2_O-to-H_2_O cycle) (Curien et al., 2016). In the Mehler reaction (Mehler, 1951), for example, O_2_ is directly reduced on the acceptor side of PSI (Kozuleva et al., 2021) to superoxide (O_2_^−^), followed by enzymatic conversion of the O_2^−^_ radical to H_2_O_2_ and O_2_ by superoxide dismutase, followed by conversion of H_2_O_2_ to H_2_O by catalase (Asada, 2000). However, if these ROS cannot be rapidly detoxified, they may damage the photosynthetic apparatus and inhibit other cellular processes. An additional reaction that facilitates non-destructive O_2_ reduction (without ROS production) involves PTOX (plastoquinol:oxygen oxidoreductase; **Figure 1A**). This enzyme controls the redox state of the plastoquinone (PQ) pool and the chloroplast stroma through chlororespiration, a light-independent process that involves the NDH complex in plants and the NDA2 protein in green algae (both function as NADPH:O_2_ oxidoreductases) (Nawrocki et al., 2015; Nixon, 2000; Peltier and Cournac, 2002; Bennoun, 1982). Moreover, when cells are exposed to light and the excitation pressure is higher than the energetic demands for CO_2_ fixation, PTOX may act as a safety valve by catalyzing a H_2_O-to-H_2_O cycle, a reaction that frequently occurs during rapid changes in the stromal redox state (e.g., transitions from dark to light or from low to high light) and during periods of environmental stress (Saroussi et al., 2016; Bailey et al., 2008; Krieger-Liszkay and Feilke, 2016; Saroussi et al., 2019). Other stromal enzymes that facilitate a H_2_O-to-H_2_O cycle include flavodiiron proteins (FLV, NADPH:flavin oxidoreductase; homologs have not been reported in angiosperms; **Figure 1A**), which oxidize NADPH generated by PSI, thereby protecting PSI (and consequently PSII) from oxidative damage during a rapid increase in reducing power that cannot be used by the CBBC (e.g., when CBBC enzymes are not fully active) (Gurrieri et al., 2021; Chaux et al., 2017; Allahverdiyeva et al., 2013; Gerotto et al., 2016).

When ROS causes damage to PSII, it can be rapidly repaired [extensively reviewed (Theis and Schroda, 2016; Komenda et al., 2012; Nath et al., 2013)], while, in contrast, recovery of PSI from oxidative damage is much slower, and in some cases irreversible (Sonoike, 2011; Heinnickel et al., 2016; Shimakawa et al., 2016; Zivcak et al., 2015). PSI photoinhibition can occur when electron flow from PSII exceeds the capacity of oxidized electron acceptors downstream of PSI, including metabolites of the CBBC (i.e., an asymmetry between PET and photosynthetic C metabolism; also known as acceptor side limitation). In this case, PSI reaction centers become reduced, O_2_^−^may be generated and react with and damage PSI iron-sulfur clusters, which can lead to degradation of PSI core subunits (Trinh et al., 2021; Sonoike, 2011; Terashima et al., 1998; Asada, 2000; Sonoike, 1996); as a consequence, the level of oxidizable P700 dramatically declines (Terashima et al., 1998, 1994; Hald et al., 2008). Although some aspects of control are known, further elucidation is required to determine how photosynthetic organisms exposed to strong abiotic fluctuations that perturb the balance between PET and CO_2_ fixation and assimilation, modulate the rate of electron transport to protect PSI from photoinhibition. There is evidence that suggests that the PGR5 protein, which is involved in cyclic electron flow (CEF) around PSI, participates in protecting PSI from damage when plants are exposed to fluctuating and high light by attenuating the rate of electron transport from PSII toward PSI (Leister et al., 2021; Yamori et al., 2016; Kono et al., 2014; Munekage et al., 2002; Suorsa et al., 2012; Munekage et al., 2008; Terashima et al., 1994); however, the mechanism associated with PGR5 function remains speculative. Another recently proposed mechanism, initially reported in cyanobacteria, describes an inhibition of Q-cycle turnover which slows down electron flow from the Cyt *b*_*6*_*f* complex to PSI [named Reduction-Induced Suppression of Electron flow, RISE (Shaku et al., 2016)]. This control of Cyt *b*_*6*_*f* activity is a consequence of the development of a proton motive force (PMF) across the thylakoid membranes (Foyer et al., 2012; Tikhonov 2014, Tikhonov 2015) and contributes to photosynthetic control, the fact that intersystem electron transport is reversibly regulated at Cyt *b*_*6*_*f*. Supporting evidence of such reversible regulation of intersystem electron transport at Cyt *b*_*6*_*f* was recently summarized (Johnson and Berry, 2021).

In this study, we created an asymmetry between PET on the one hand, and growth, CO_2_ fixation and assimilation, on the other, by culturing a *Chlamydomonas reinhardtii* (designated Chlamydomonas throughout) mutant (*sta6*) that cannot synthesize starch (Zabawinski et al., 2001; Li et al., 2010) in medium devoid of nitrogen (N). This led to the establishment of a photosensitive regulatory pathway that appears to protect PSI from photodamage by modulating electron flow through the Cyt *b*_*6*_*f* complex, which may reflect photosynthetic control, while at the same time sustaining PSII activity by redirecting photosynthetic electron flow to the reduction of O_2_ through PTOX.

## RESULTS

### Asymmetry between absorbed light energy and carbon metabolism attenuates O_2_ evolution and electron transport

To disrupt the balance between the absorbed light energy and C metabolism, we deprived Chlamydomonas of N. This treatment, which resulted in the cessation of cell growth, eliminated the main cellular sink (growth) for photosynthetically generated reductant, ATP, and C backbones. N-deprived Chlamydomonas cells rapidly acclimate to limiting N conditions by activating expression of specific pathways involved in N acquisition and lipid and starch synthesis (Davey et al., 2014; Siaut et al., 2011); the synthesis of starch and triacylglycerides, N-free C polymers, serve as sinks for energy and fixed C generated by PET and the CBBC. To further eliminate potential pathways for assimilating and storing fixed C and utilizing ATP, we combined N deprivation with the use of the Chlamydomonas *sta6* mutant (Zabawinski et al., 2001). This mutant is null for the small subunit of ADP-glucose pyrophosphorylase (AGPase), an enzyme that catalyzes the synthesis of ADP-glucose from glucose-1-phosphate and ATP (**Figure 1B**), the rate-limiting step in starch synthesis. The *sta6* mutant, which makes no detectable starch (**Figure S1)**, can be phenotypically rescued with an ectopically expressed, wild type (WT) copy of the *STA6* gene; the rescued strain, C6 (*sta6::STA6*), synthesizes starch; (**Figure S1)** (Li et al., 2010). The genome of the C6 strain was recently sequenced and has been used as a reference for evaluating the *sta6* phenotype (Siaut et al., 2011; Schmollinger et al., 2014; Blaby et al., 2013; Anderson et al., 2016). To assess the balance between PET and downstream C metabolism in *sta6* and C6, we cultured Chlamydomonas cells for 24 h at 100 µmol photons m^−2^s^−1^ in MOPS-Tris medium (see METHODS) devoid of N and then measured net O_2_ evolution immediately following a dark-to-light transition (experimental design in **Figure 1C**). N-deprived C6 maintained high rates of net O_2_ evolution (Pmax_net_ = ∼160 μmol O_2_ mg Chl^−1^ h^−1^, black curve in **Figure 1D**), similar to those of C6 cultured in the presence of N (∼180 μmol O_2_ mg Chl^−1^ h^−1^, **Figure S2**). In contrast, net O_2_ evolution in the N-deprived *sta6* mutant was nearly undetectable (i.e., near the O_2_ compensation point; **Figure 1D**) under N deprivation conditions, while in the presence of N, net O_2_ evolution was ∼60 μmol O_2_ mg Chl^−1^ h^−1^ (**Figure S2**). These results suggest that a depressed ability to use reductant because of the loss of two major electron sinks resulted in strong disruption of the balance between PET and downstream photosynthetic C metabolism.

To further characterize PET with respect to C metabolism, we monitored linear electron flow (LEF) in *sta6* and C6 using fluorescence spectroscopy. Following a period of 24 h of N deprivation, the effective PSII quantum yield (ΦPSII) of *sta6* was much lower than that of C6 at all tested light intensities, while no strong differences in ΦPSII between the strains were observed when the cells were cultured in the presence of N (**Figure S3**). We also measured the re-reduction rate of P700 in the presence of DCMU and hydroxylamine to evaluate the contribution of NDA2-dependent reduction of P700 (Saroussi et al., 2016). The rate of P700 re-reduction in N-deprived, WT Chlamydomonas was previously shown to increase as a consequence of elevation of the NDA2-dependent pathway (Saroussi et al., 2016). In this pathway, electrons derived from NADPH are used to reduce the PQ pool. In the absence of N, the re-reduction rate of P700 measured for C6 increased by ∼1.5 fold (relative to the re-reduction rate in the presence of N; **Figure S4A** and **B**), similar to the observation previously made for WT Chlamydomonas cells (Saroussi et al., 2016). However, unlike C6, N deprivation of *sta6* resulted in a slowing of the rate of P700 re-reduction (**Figure S4A** and **B**). Hence, N-deprived *sta6* has both diminished LEF and NDA2-dependent re-reduction of PSI.

We also explored the possibility that photosynthetically generated electrons in N-deprived C6 and *sta6* were shuttled to the mitochondrial respiratory electron transport chain. N-deprived and N-replete C6 displayed similar rates of dark respiration immediately following the extinction of the light (**Figure S5A**). In contrast, while respiratory O_2_ uptake was observed in *sta6* grown in the presence of N, almost no dark respiration was observed in N-deprived *sta6* (**Figure S5A** and **B**). In sum, in the absence of starch synthesis (*sta*6) during N deprivation, PET, net O_2_ evolution, and mitochondrial respiration were all severely diminished.

The negligible net O_2_ production exhibited by N-deprived *sta6* cells following a dark-to-light transition (**Figure 1D**) can be explained by one or a combination of the following: **a**. Feedback on electron transport caused by the extreme imbalance between PET and C metabolism leads to inactivation of PSII which inhibits water oxidation, O_2_ production, and electron transport; **b**. PSII remains active (i.e., H_2_O is oxidized), but the O_2_ produced is consumed through activation of a H_2_O-to-H_2_O cycle associated with alternative outlets for photosynthetic electrons (Curien et al., 2016); **c**. An electron transport reaction downstream of PSII is restricted that slows the rate of H_2_O oxidation. To evaluate these hypotheses, we first analyzed the maximal quantum efficiency of PSII (Fv/Fm) and whether the level of the PSII core subunit, D1, was affected under our experimental conditions. We observe a 20% change in the Fv/Fm of *sta6* cells deprived of N while no change was observed for C6 (comparing *sta6* with C6 cells under -N and +N conditions, **Figure S6A**), which does not necessarily reflect the extent of photoinhibition, while the levels of the PSII reaction center protein (determined immunologically) remained approximately the same (normalized to tubulin, **Figure S6B**). Furthermore, to assess PSII activity, we measured the capacity of C6 and *sta6* cells to evolve O_2_ in the presence of *p*-benzoquinone (PBQ), an artificial electron acceptor that oxidizes the PQ pool. The rate of O_2_ evolution in the presence of this compound reflects the maximal capacity of PSII to oxidize H_2_O and evolve O_2_. No significant differences in the levels of O2 evolution were observed with the addition of PBQ to C6 and *sta6* cells that were cultured in the presence of N (**Figure S6C** and **D**). Under N deprivation conditions, however, the net rate of O_2_ evolution in *sta6* cells was zero (similar results in **Figure 1D**); most of the activity was regained when the cells were supplemented with PBQ. These results demonstrate that PSII of N-deprived *sta6* is still active.

We next monitored the real-time exchange of O_2_ in N-deprived *sta6* cells using membrane inlet mass spectrometry (MIMS) to distinguish between light-dependent production and uptake of O_2_ (monitoring the ^16^O_2_ signal for production and the ^18^O_2_ signal for uptake; **Figure 2**). N-deprived *sta6*, treated as indicated in **Figure 1C** (i.e., dark-to-light transition), produced O_2_ at a maximal rate (Pmax) of ∼25 µmol O_2_ mg Chl^−1^ h^−1^ soon after being exposed to 300 µmol photons m^−2^ s^−1^ (**Figure 2A**). However, the rate of O_2_ production and uptake were similar (compare **Figure 2A** with **2B**), which resulted in approximately zero net O_2_ evolution (**Figures 2C** and **Figure 1D**). Net zero O_2_ evolution was not observed when *sta6* cells were cultured in the presence of N (compare **Figures 2** and **S7A-C**) or when identical analyses were performed with C6 cells. C6 cells displayed a high rate of net O_2_ production with similar kinetics for O_2_ production/uptake regardless of the N status of the cells (**Figure S8A-G**; and as shown in (Saroussi et al., 2019)), although the absolute level of production and uptake were slightly lower for N-deprived C6. These results suggest that diminished C metabolism in the N-deprived *sta6* mutant resulted in a marked reduction in the rate of gross O_2_ evolution and activation of alternative terminal electron outlets that consume all of the evolved O_2_ (and electrons) generated by PSII-dependent H_2_O oxidation.

**Figure 2:**
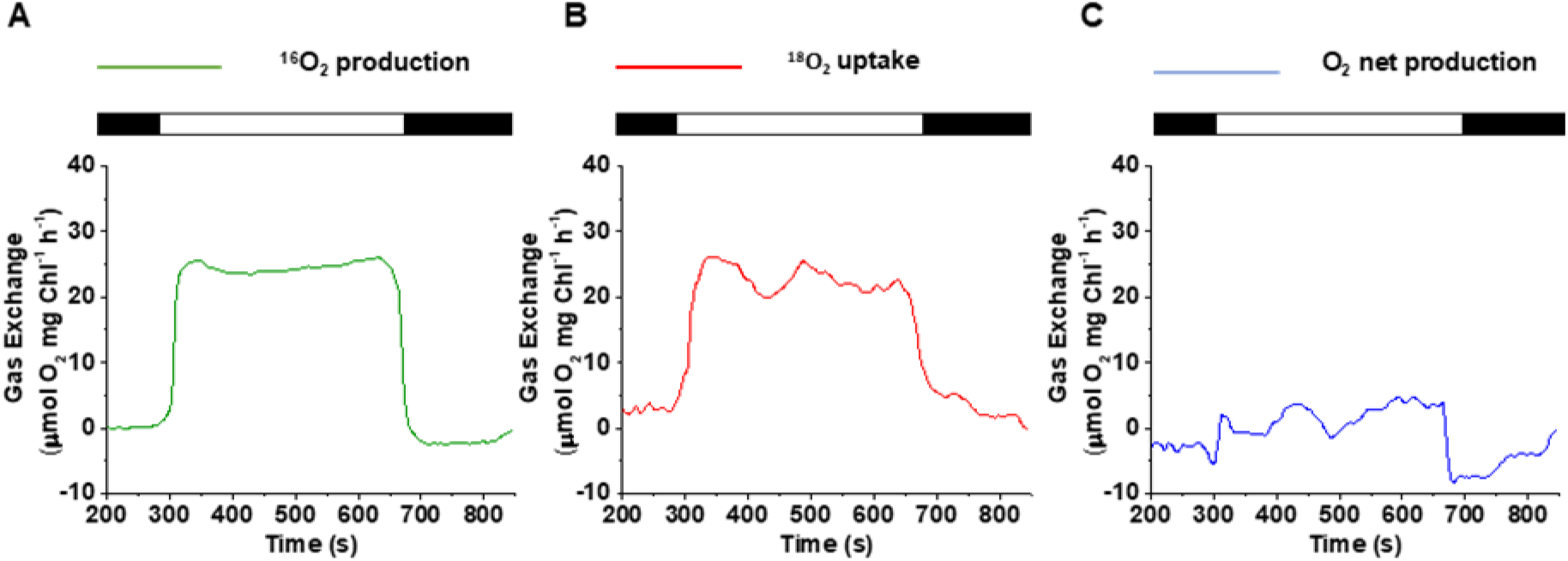
Real-time O_2_ exchange for the N-deprived *sta6* mutant using MIMS: **A**. Light-induced gross O_2_ production (^16^O_2_ evolution, green curve), and **B**. uptake (^18^O_2_ consumption, red curve), for N-deprived *sta6* following a dark (30 min) to light (300 μmol photons m^−2^ s^−1^) transition; gas exchange was monitored for 6 min in the light using MIMS. Prior to turning on the light, the cells were maintained for 5 min in the dark (in the MIMS chamber) and then for 3 min in the dark following illumination. **C**. Net O_2_ production (blue curve), was calculated according to the formula given in the **Methods**. Each trace represents an average of at least 3 biological replicates ± SE.

### N-deprived *sta6* cells have a highly reduced quinone pool

We next explored the mechanism underlying impaired O_2_ production and electron transport in the *sta6* mutant (**Figures 2, S3** and **S4**). To determine if the pool of electron acceptors downstream of PSII were sufficient to support PSII activity, we quantified the photochemical efficiency (qL) of N-deprived C6 and *sta6* as a function of light intensity (**Figure 3**). The photosynthetic parameter qL indicates the redox state of Q_A_, the primary electron acceptor of the PSII reaction center. Assuming that Q_A_ and Q_B_ are in equilibrium, qL would reflect the redox status of the PQ pool; a lower qL value indicates a more reduced electron transport chain (Kramer et al., 2004). To create a direct relationship between the qL value and the redox state, we used an inverse function, 1-qL, which positively correlates with the PQ pool redox state. Following dark acclimation, the 1-qL values for *sta6* cells were higher at any given light intensity than the 1-qL values measured for C6 cells, suggesting a more negative redox potential (more reduced PQ pool) in *sta6* relative to C6 at each of the light intensities. Even when exposed to low light (LL, 15 µmol photons m^−2^ s^−1^), the 1-qL value was at 70% maximum for *sta6* (suggesting a highly reduced PQ pool). In contrast, for C6 at the same light intensity, the 1-qL value attained only 20% of the maximum value (**Figure 3A**). Surprisingly, following a 1 h light acclimation of *sta6*, the 1-qL levels declined and were only slightly higher than those of C6; that is, the 1-qL values following exposure of *sta6* to LL indicated a much more highly oxidized PQ pool (**Figure 3B**, compare red solid and dashed curves). After exposure to the same light conditions, we did not observe a significant change in 1-qL values for C6 (compare black curves in **Figure 3A** and **3B**). These results suggest that following a period of dark acclimation, the PQ pool of the N-deprived *sta6* mutant is maintained in a highly reduced state, although the redox pressure can be significantly relieved by a 1 h exposure to light. These observations led to the hypothesis that N-deprived *sta6* experiences a reversible restriction in PET downstream of the PQ pool.

**Figure 3:**
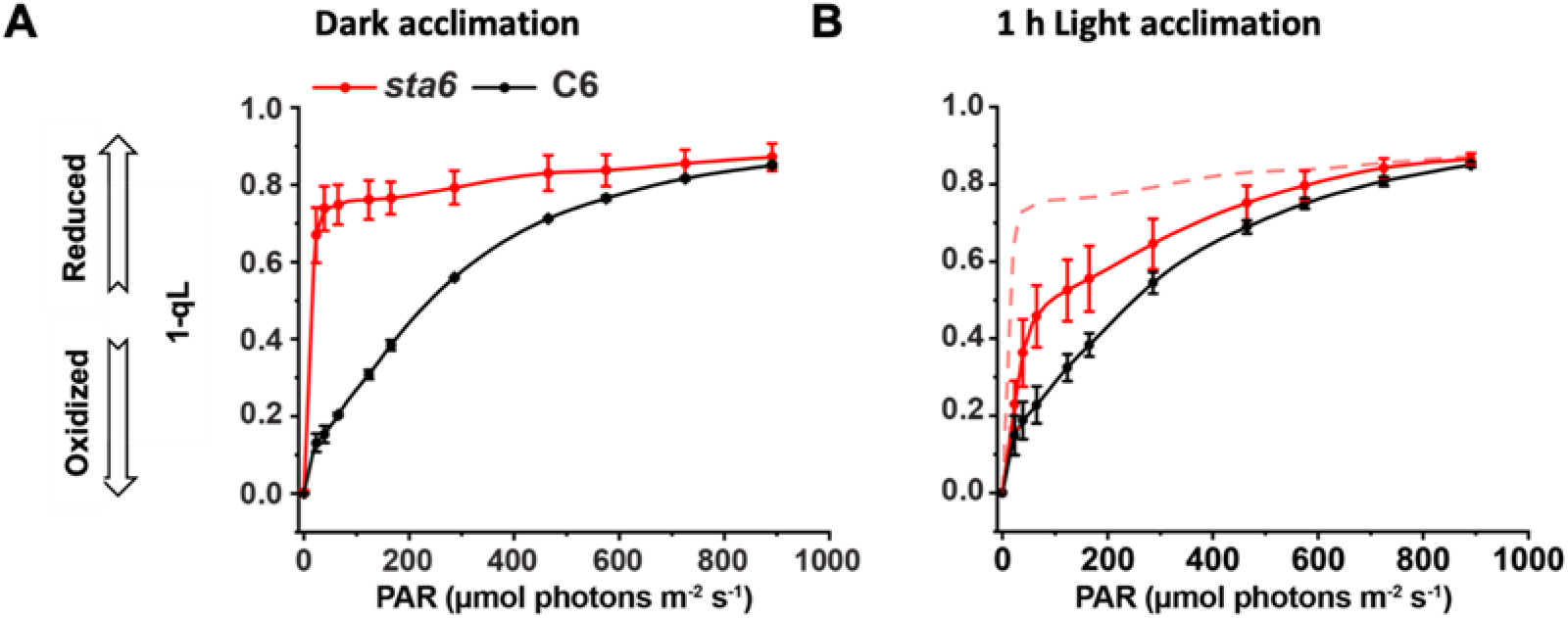
Redox state of the PQ pool: 1-qL, used to estimate the redox state of the PQ pool, was calculated for C6 and *sta6* (black and red curves, respectively), from a light curve in which the fluorescence output was quantified using a Dual-PAM fluorometer (Walz, Germany) following **A**, dark acclimation, and **B**, 1 h of acclimation to 100 µmol photons m^−2^ s^−1^. For comparison, we included the 1-qL data of dark-acclimated *sta6*, which is shown as the dashed red line in panel **B**. Each data point reflects an average of at least 3 biological replicates ± SD.

### Cyt *b*_*6*_*f* complex is the site of a reversible restriction in electron transport

The hyper-reduced PQ pool in dark-acclimated, N-deprived *sta6* suggested a downstream limitation in electron flow that impairs quinol oxidation. To explore this issue, we measured the oxidation/reduction kinetics of Cyt *f* in N-replete and deprived *sta6* cells; Cyt *f* oxidation reflects electron flow from Cyt *f* to PC while re-reduction reflects electron flow from the iron-sulfur cluster of the Rieske iron-sulfur protein (ISP) to Cyt *f*. Initially, *sta6* cells were cultured in the presence of N. After the cells were dark acclimated, they were exposed to a laser flash that oxidized Cyt *f*, which was then re-reduced with a half time (T_1/2_) of 24 ms (black curve of **Figure 4A** and black boxplot in **Figure 4C**). However, if the cells were light acclimated (put back under standard culture conditions at 100 µmol photons m^−2^ s^−1^ for 10 min prior to the measurements), the T_1/2_ for re-reduction of Cyt *f* was 5 ms (red curve in **Figure 4A** and red boxplot in **Figure 4C**); this value is similar to the reported re-reduction rate of Cyt *f* in WT Chlamydomonas cells (de Vitry et al., 2004). A similar trend for Cyt *f* re-reduction kinetics was observed when N-replete C6 cells were subjected to the same dark-to-light transition (16 ms and 7 ms for dark- and 10 min light-acclimated cells, respectively; **Inset** of **Figure 4A**). Surprisingly, repeating the same analyses with dark-acclimated *sta6* cells cultured in a medium devoid of N yielded a T_1/2_ re-reduction rate for Cyt *f* of 66 ms, which is much slower than in the presence of N (compare black curves in **Figures 4B** and **Figure 4A** and black boxplot in **Figure 4D** with **4C**). However, after exposing N-deprived, dark-acclimated *sta6* to light (same as above) for 10 min, the T_1/2_ for re-reduction of Cyt *f* was reduced to 28 ms (red curve of **Figures 4B** and red boxplot in **Figure 4D**) while after 1 h of light exposure the T_1/2_ for re-reduction was 17 ms (blue curve of **Figures 4B** and blue boxplot in **Figure 4D**). While the N-deprived C6 cells also exhibited some slowing of Cyt *f* re-reduction kinetics after dark acclimation (T_1/2_ = 39 ms) (**Inset** in **Figure 4B)**, unlike *sta6* cells, re-reduction of Cyt *f* was fast (similar to WT nutrient replete) following exposure of the cells to just 10 min of light (3 ms, **Inset** of **Figure 4B)**. This modulated flow of electrons to Cyt *f* reflects a reversible restriction of electron transport that appears to be triggered by environmental conditions.

**Figure 4:**
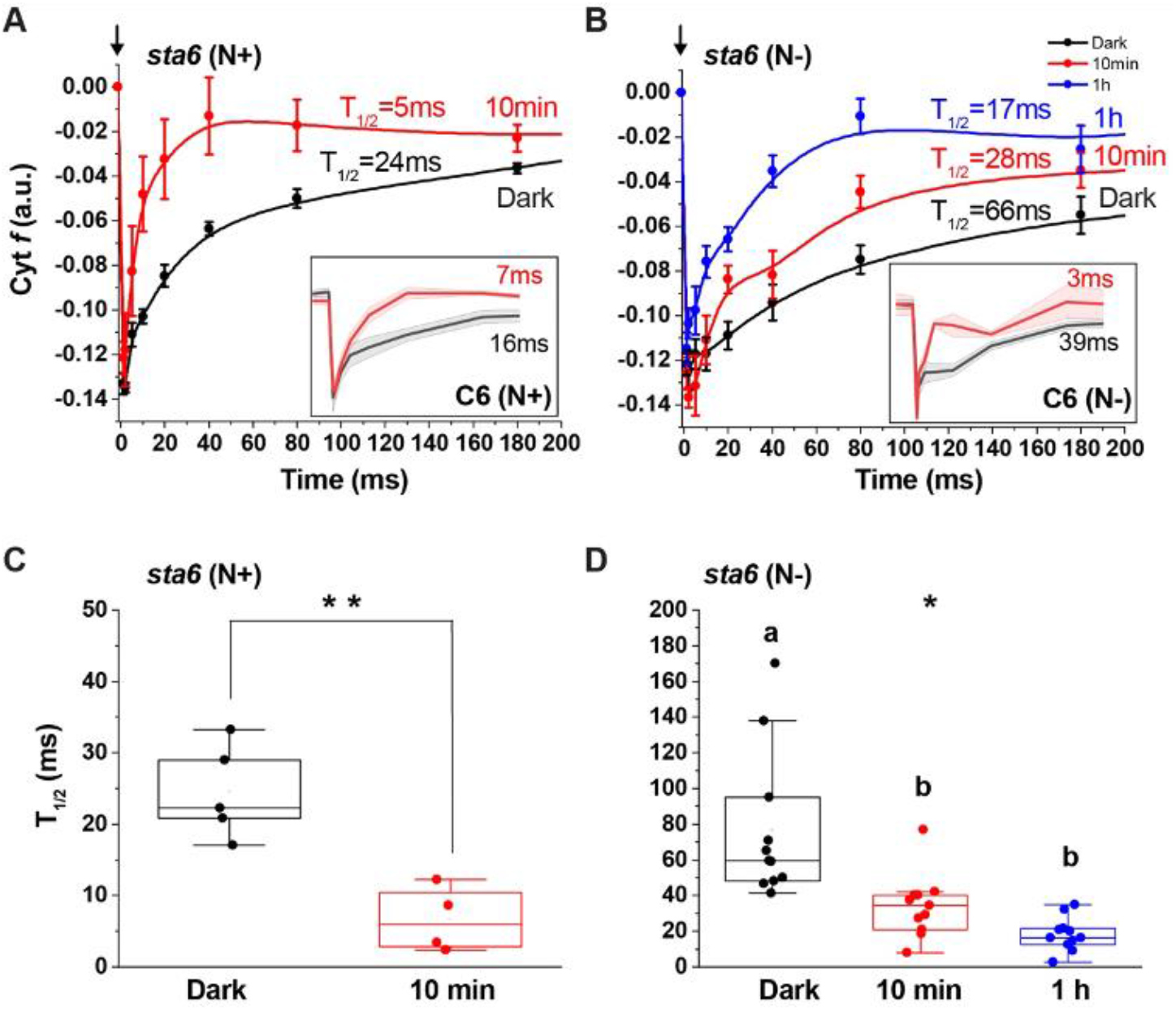
Reversible restriction of electron transport to Cyt *f*: Re-reduction kinetics of Cyt *f* in dark- and light-acclimated (100 µmol photons m^−2^ s^−1^) *sta6* cultured in the presence and absence of N (**A** and **B**, respectively). A saturating laser flash (black arrow) was administered at time 0, immediately following the dark incubation, to oxidize Cyt *f*. Re-reduction kinetics were monitored by absorbance changes (described in **Methods**) at the indicated times following the flash. Half time (T_1/2_) of re-reduction was calculated by fitting the re-reduction component of Cyt *f* kinetics to a first order decay equation (although the kinetics may be more complex). All *p*-values for these fittings were <0.05. The data are an average of at least five biological replicates ±SE. **Inset:** Cyt *f* oxidation reduction kinetics of N-replete (**A**) and N-deprived (**B**) C6 cells. For N-replete C6 cells (left panel) the T_1/2_ for Cyt *f* re-reduction are 16 ± 3.2 and 7 ms ± 0.9 for the dark- and light-acclimated (10 min) cells, respectively. For N-deprived C6 cells (right panel) the T_1/2_ for Cyt *f* re-reduction are 39 ± 4 and 3 ms ± 0.6 for the dark- and light-acclimated (10 min) cells, respectively. Panels **C** and **D** show preset boxplots of the Cyt *f* T_1/2_ re-reduction rates following the dark incubation, or after 10 min and 1 h in the light following the dark incubation. For **C**, t-test ** *p*-value < 0.005. For **D**, One-way ANOVA, * *P-*value < 0.05; Post-hoc Tukey, *p*-value < 0.05. Note the different scales of the y-axes of panels **C** and **D** (there is a significant difference in the T_1/2_ values following 10 min of light exposure; t-test, *p-*value =0.0018).

We also sought to determine if the reversible restriction in electron transport was a function of an imbalance between the amounts of Cyt *f* and PC in the N-deprived *sta6* cells. Based on immunodetection, the accumulation of Cyt *f* and PC were similar in dark- and light-acclimated, N-deprived *sta6* cells. There was also no difference in the accumulation of these proteins in N-deprived, dark- and light-acclimated C6 cells, although the levels of both Cyt *f* and PC appeared to be slightly lower in the mutant than in the complemented strain (**Figure S9**). Overall, these results suggest that the difference between electron transport in dark- and light-acclimated cells was not affected by changes in the levels of these proteins.

Lastly, we explored the possibility that a low ATP turnover rate in *sta6* leads to a compromised chloroplast ATP synthase activity which feeds back on PET (slowing PET) to increase the reduction state of the PQ pool. To test this hypothesis, we repeated the 1-qL measurements shown in **Figure 3** for N-deprived *sta6* in the presence of a protonophore and ionophore. We used nigericin, a drug that dissipates the H^+^ gradient across the thylakoid membranes, which would eliminate the feedback of ATP synthesis on PET, and valinomycin, a drug that dissipates the electrochemical gradient across the thylakoid membranes. Upon supplementing the cells with 10 µM nigericin, 10 µM valinomycin, or both, 1-qL remained the same as that of non-treated cells (**Figure S10**), suggesting that PET was not restricted as a consequence of the rate of ATP synthesis. Hence, the H^+^ and electrochemical gradients across the thylakoid membranes do not appear to be required to sustain a restriction in PET across the Cyt *b*_*6*_*f* complex.

### Blocking electron flow sustains oxidized P700 and photoprotection

To validate the reversible restriction in electron transport at the level of Cyt *b*_*6*_*f*, we analyzed the redox state of P700. Diminished electron flow out of Cyt *b*_*6*_*f* would result in a mostly oxidized P700 following exposure of the cells to light. As the restriction is removed, the P700 redox status should become similar to that of cells in which no or little restriction was experienced. To examine this, we monitored oxidation/reduction kinetics of P700 in N-deprived *sta6* and C6 cells (**Figure 5**) that were treated identically to those cells used for the experiment given in **Figure 4**. Importantly, the cells were not treated with DCMU, a drug that inhibits electron flow out of PSII, to allow electrons originating from both LEF and CEF to reduce oxidized P700. Following illumination of dark-acclimated C6 cells with non-saturating light (60 µmol photons m^−2^ s^−1^ for 10 sec), ∼25% of the P700 was oxidized at the end of the light period (black curve, left panels of **Figures 5A** and left bar graph of **Figure 5B**). In contrast, for N-deprived, dark acclimated *sta6* cells, nearly 80% of the P700 remained oxidized at the end of the actinic light period (red curve, left panel of **Figures 5A** and left bar graph of **Figure 5B**), suggesting that electron flow from Cyt *b*_*6*_*f* to PSI was restricted in N-deprived *sta6* cells immediately following the dark period; this restriction would potentially prevent (or diminish) PSI photoinhibition when the cells are limited for downstream electron acceptors. We then exposed N-deprived *sta6* to 100 µmol photons m^−2^ s^− 1^ for 1 h (similar to 1 h treatment of cells used in the measurements of 1-qL and Cyt *f* re-reduction kinetics; **Figures 3B** and **4B**). Following this light acclimation period, P700 oxidation/reduction kinetics in C6 and *sta6* were similar, with the proportion of oxidized P700 just over 25% for both strains at the end of the actinic light exposure (right panel in **Figures 5A** and right bar graph of **Figure 5B**), supporting the conclusion that N-deprived *sta6* cells experience a reversible restriction in electron flow to Cyt *f*. Moreover, the total amount of photo-oxidized P700 was identical for the two strains (t-test, *p*-value = 0.66), suggesting that PSI in *sta6* did not experience more oxidative damage than in C6 during the dark-to-light transition, despite the inability of *sta6* to store starch, a major sink for assimilating electrons downstream of PSI during N deprivation.

**Figure 5:**
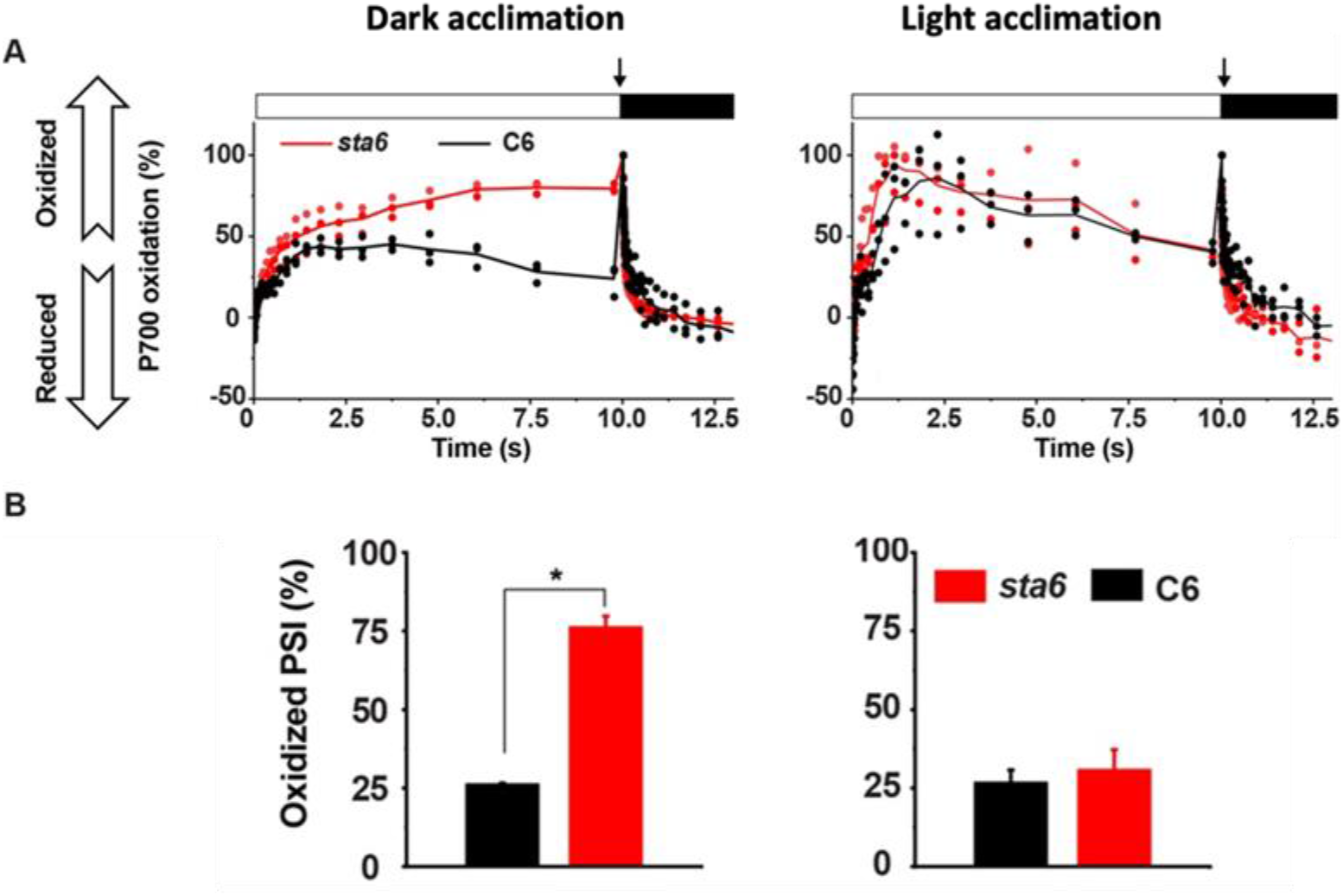
The impact of light on the oxidation state of P700. **A**. P700 oxidation/reduction kinetics of N-deprived C6 and *sta6* following dark acclimation (left), and light acclimation for 1 h (100 µmol photons m^−2^ s^−1^), (right). A saturating flash (black arrow) was administered at the end of the actinic light exposure to fully oxidize P700. Values for all biological replicates (N=3) are presented. **B**. Levels of P700^+^ (oxidized PSI) in dark-acclimated (left) and light-acclimated (right) C6 and *sta6* cells. Data was calculated from the kinetics shown in Panel **A**. * t-test, *p*-value = 0.0013; N=3 ± SD. 3 biological replicates ± SE.

### PTOX-dependent H_2_O-to-H_2_O cycle drives PSII activity

While PSII was still functioning in N-deprived *sta6* following a dark-to-light transition (Fv/Fm was >0.5, **Figures 2** and **S6**), net O_2_ production was approximately zero (**Figures 1D** and **2C**) because the amount of O_2_ released as a consequence of H_2_O oxidation was essentially identical to the amount of O_2_ consumed (**Figure 2**). Given the restriction in electron flow at the level of Cyt *b*_*6*_*f* (**Figure 4**), the most plausible alternative outlet that could facilitate reduction of O_2_ to H_2_O (i.e., O_2_ consumption) upstream of Cyt *b*_*6*_*f* in the electron transport chain is the alternative oxidase PTOX, which oxidizes a quinol molecule and reduces O_2_ to H_2_O (Nawrocki et al., 2015) (**Figure 6A**). To evaluate this possibility, we repeated the MIMS experiment shown in **Figure 2**, but in the presence of 2 mM propyl gallate (PG), a drug that inhibits alternative oxidases such as PTOX. Following dark acclimation (30 min, as in **Figure 1C**), the rates of light-dependent O_2_ production and uptake in N-deprived *sta6* were dramatically diminished (300 µmol photons m^−2^ s^−1^; solid green and red curves in **Figure 6B**; note the y axis), with PG-treated *sta6* reaching a Pmax 15-fold more slowly than the untreated cells (T_1/2_ = ∼ 84 s for PG-treated and ∼ 5 s for untreated cells; green bars in **Figure 6C** and compare solid and dashed green curves in **Figure 6B)**. The O_2_ uptake rate for PG-treated *sta6* was also diminished, reaching maximal uptake (Umax) 10-fold more slowly than the untreated cells (T_1/2_ = ∼ 59 s for PG treated and ∼ 5 s for untreated cells; red bars in **Figure 6C** and compare solid and dashed red curves in **Figure 6B**). Importantly, O_2_ production and uptake were directly correlated over the light period and the rate at which Umax and Pmax were obtained were not significantly different (59 ± 7 s, and 84 ± 28 s, respectively; N=3; t-test, *p-*value = 0.26; compare green and red bars in **Figure 6C** and green and red curves in **Figure 6B**); as a result, net O_2_ production was close to zero (blue curve, **Figure 6B**). We note that as O_2_ uptake in the light proceeds, the effect of PG diminishes (>100 sec), which may be a consequence of light inactivation of the inhibitor or a loss of the restriction of electron flow through Cyt *b*_*6*_*f* because PTOX activity is critical for sustaining the restriction. In addition, we examined the effect of PG on electron transport in N-deprived *sta6* cells (**Figure S11**). While the restriction of electron transport across the Cyt *b*_*6*_*f* complex was not affected by PG (the switch remains restrictive, **Figure S11A**), electron flow generated by PSII (i.e.,ΦPSII) further decreased in low/moderate light (**Figure S11B**), congruent with the observation that O_2_ production is diminished when PTOX is inhibited (**Figure 6B**). In contrast, inactivation of PTOX in N-deprived C6 cells did not significantly impact the rate of O_2_ production or uptake (Pmax = ∼120 µmol O_2_ mg Chl^−1^ h^−1^ for both treated and untreated cells; T_1/2_ to obtain Pmax for treated and untreated cells was 19.4 ± 2.1 s and 15.9 ± 0.23 s, respectively; **Figure 6D**). Together, these data indicate that following a restriction in electron flow at the Cyt *b*_*6*_*f* complex in the N-deprived *sta6* mutant, inactivation of PTOX by PG strongly inhibits both the production and uptake of O_2_; PG does not affect O_2_ production in C6. Therefore, the H_2_O-to-H_2_O cycle catalyzed by PTOX appears to be the driving force for O_2_ production in the N-deprived *sta6* mutant, supporting the conclusion that there is a reversible restriction of electron flow across the Cyt *b*_*6*_*f* complex in N-deprived *sta6* cells during a dark-to-light transition.

**Figure 6:**
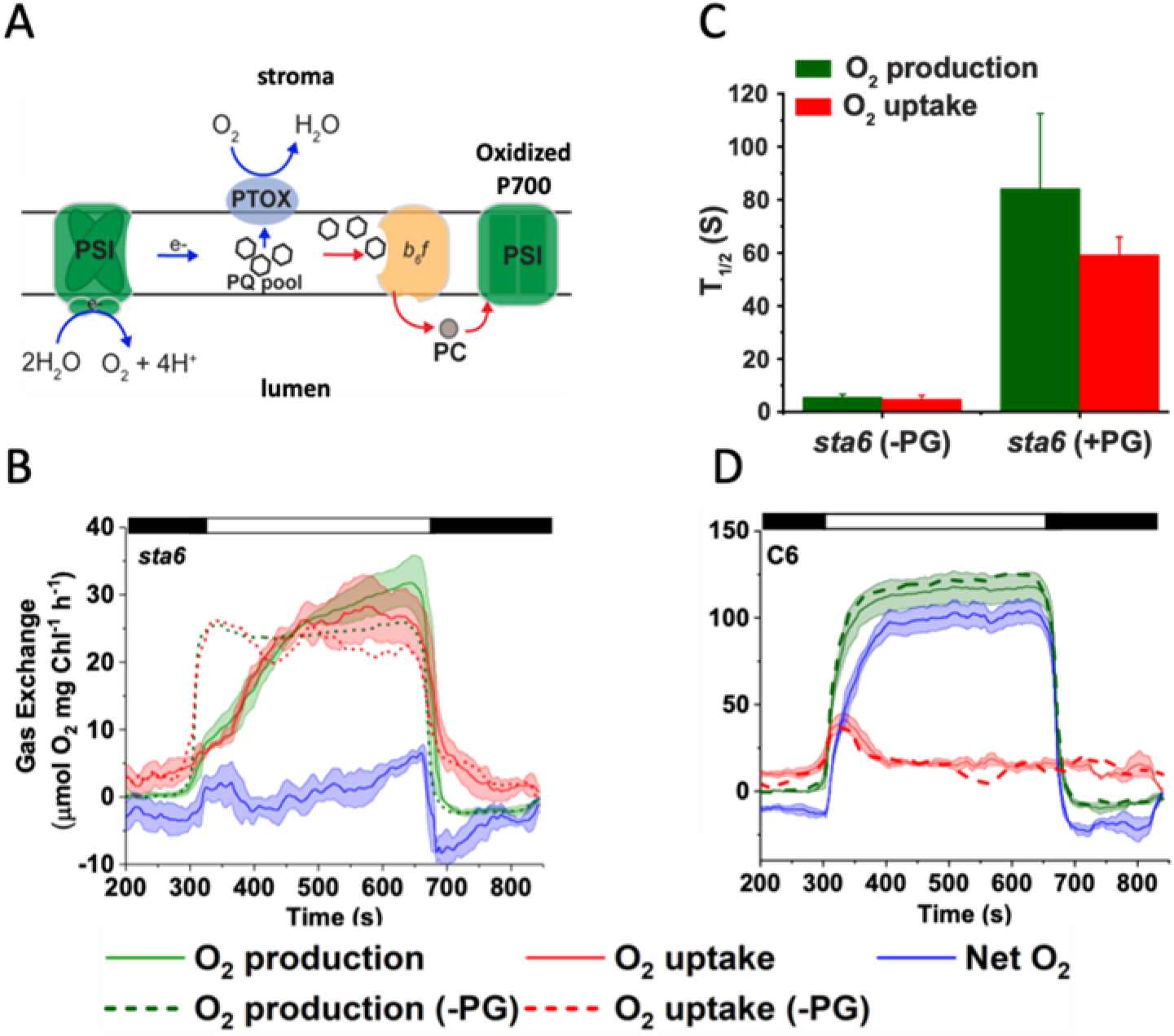
Real-time O_2_ exchange for N-deprived cells in the presence of a PTOX inhibitor: **A**. Schematic of photosynthetic electron flow when electron flow across the Cyt *b*_*6*_*f* complex is restricted in N-deprived *sta6*. Red and blue arrows indicate restricted and non-restricted pathways, respectively. **B**. Gross O_2_ production (^16^O_2_ evolution, green curves) and gross O_2_ uptake (^18^O_2_ consumption, red curves) are presented for N-deprived *sta6* both in the presence (solid curves) and absence (broken curves) of 2 mM propyl gallate (PG), an inhibitor of PTOX. Net O_2_ production in the presence of PG is shown by the blue curves. Experimental design was as in **Figure 2**. Each curve represents an average of at least 3 biological replicates ± SE. **C**. T_1/2_ for attaining Pmax (green bars) and Umax (red bars) in N-deprived *sta6* in the presence and absence of PG. Rates were calculated after fitting the O_2_ production and uptake curves to a first order exponential decay. The presented data is an average of at least 3 biological replicates ± SD. **D**. Gross O_2_ production (^16^O_2_ evolution, green curves) and gross O_2_ uptake (^18^O_2_ consumption, red curves) are presented for N-deprived C6 both in the presence (solid curves) and absence (broken curves) of 2 mM PG. Net O_2_ production in the presence of PG is shown by the blue curve. Experimental design was as in **Figure 2**. Each curve represents an average of at least 3 biological replicates ± SE.

## DISCUSSION

Abiotic conditions on Earth, including temperature, light, moisture, and levels of available nutrients, are rapidly being altered in response to global climate change. Changing environmental conditions can constrain the growth and development of photosynthetic organisms and lead to an imbalance between the light energy absorbed by the photosynthetic apparatus and its use for the assimilation of CO_2_ by the CBBC, growth and the storage of starch and lipids. When such imbalances occur, some of the solar energy absorbed by PSI and PSII can be converted to ROS which creates oxidative stress and consequently photoinhibition, degradation of photosynthetic reaction centers, and damage to essential cellular components such as nucleic acids, proteins, and lipids. While an efficient system has evolved to repair damaged PSII, there is no corresponding system associated with PSI; recovery of PSI is slow or irreversible, which can strongly impact plant primary productivity. Hence, photosynthetic organisms have evolved mechanisms to dissipate excess absorbed light energy through NPQ (Müller et al., 2001) and the use of shunts and alternative electron valves (not directly involved in CO_2_ fixation) for photochemical quenching, many of which consume O_2_ (Saroussi et al., 2019; Curien et al., 2016; Roach and Krieger-Liszkay, 2014), and the implementation of photosynthetic control, which alters the conductance of electrons through the Cyt *b*_*6*_*f* complex, similar to a transistor as it was previously described (Johnson and Berry, 2021. The integration of these processes favor optimizing the use of light energy for CO_2_ fixation, while at the same time protecting reaction centers from damage. In this work, we explored mechanisms by which Chlamydomonas protects PSI from photoinhibition when there is marked asymmetry between the rate at which energy is absorbed and can be used in C metabolism.

Chlamydomonas cells exposed to N deprivation undergo rapid metabolic acclimation. The cells stop growing and the light-generated reductant and ATP drive assimilation of CO_2_, mostly into starch and lipids (Davey et al., 2014; Siaut et al., 2011; Smith and Gilmour 2018). It is not surprising that a severe imbalance between solar energy absorption by reaction centers and C assimilation occurs when Chlamydomonas cells are both deprived of N (limits Ci utilization for growth) and are unable to synthesize starch, which is the case for the *sta6* mutant; the imbalance would likely be most severe during dark-to-light transitions, conditions in which the CBBC is not initially fully active (Buchanan, 2016; Michelet et al., 2013; Wirtz et al., 1982; Saroussi et al., 2019) (**Figure 1A**). Consequently, the pool of available electron acceptors downstream of PSI would be severely limited, potentially leading to ROS accumulation and the irreversible photoinhibition of PSI (Asada, 2000; Allahverdiyeva et al., 2015; Sonoike, 1996, 2011).

A detailed analysis of electron transport in N-deprived *sta6* cells demonstrated that following dark acclimation, Cyt *f* was readily oxidized by a laser flash (normal Cyt *f* oxidation; **Figure 4B**), but that its rate of re-reduction was very slow relative to WT cells or the complemented strain. These results indicated a restriction in electron flow between the PQ pool (or Q_o_) and Cyt *f*, raising the possibility that the restriction is located between the Rieske protein and Cyt *f* (Sarewicz et al., 2017). In accord with this possibility, the mutant exhibited a marked reduction in the rate of quinol oxidation, as indicated by hyper-reduction of the PQ pool (**Figure 3A**) and diminished rates of LEF, P700 re-reduction via the NDA2 pathway, and mitochondrial respiration (**Figures S3-S5**). It was previously shown that when oxidation processes downstream of Cyt *b*_*6*_*f* are arrested or slowed, the iron-sulfur cluster of the complex adopts a metastable state that maintains electrons at a local energetic minimum making them nonreactive with O_2_ (Sarewicz et al., 2017). This stable cluster state of the Cyt *b*_*6*_*f* complex can sustain the block in electron transport without generating ROS, which might reflect altered interactions or accessibility between the iron-sulfur clusters and O_2_.

The electron transport restriction in dark acclimated, N-deprived *sta6* cells resulted in elevated light-dependent P700 oxidization (i.e., P700 mostly in the P700^+^ state; left panels of **Figures 5A** and **5B**), a state in which PSI is photo-protected since P700^+^ is an efficient quencher of absorbed excitation energy (i.e., converts it to heat) (Shubin et al., 2008), and the oxidized P700 cannot readily generate O_2_^−^ because of its positive charge. The ‘restricted state’ could be reversed upon exposure of the dark-acclimated, N-deprived *sta6* cells to sub-saturating light. Interestingly, the restricted state could be reversed, resulting in low levels of O_2_ evolution following exposure of the dark-acclimated, N-deprived *sta6* cells to sub-saturating light (**Figures 3B** and **4B**); the time required for reversal in the light requires more than 10 min and is nearly complete in 1 h. These kinetic features may indicate a biochemical modification of the Cyt *b*_*6*_*f* complex, potentially post-translational modification(s) and or activation of a pathway that consumes these electrons. Such a pathway might include the synthesis of lipids, which typically occurs in N deprived Chlamydomonas cells (Xu et al., 2008, Siaut et al., 2011), although this would need to be examined in longer term experiments (e.g. transfer to light reversal conditions for several hours instead of just one hour). Additionally, after this light reversal, the kinetics of P700 oxidation/reduction in N-deprived *sta6* and C6 were similar (N-deprived C6 does not exhibit a similarly severe restriction in electron transport after incubation in the dark), suggesting that Cyt *b*_*6*_*f* associated regulation of PET may protect PSI in N-deprived *sta6* from oxidative damage as the cells transition to the light. The regulation of electron conductivity between PSII and PSI according to the cell’s capacity to utilize reducing equivalents and ATP for CO_2_ fixation and in C assimilatory pathways (while protecting PSI) has been designated ‘photosynthetic control’ (Foyer et al., 1990; Weis et al., 1987). The specific factors that govern photosynthetic control have not been fully detailed, although studies are providing information concerning its mechanistic features and potentially its complexity. As an example, PSI photoinhibition was analyzed in two *Nicotiana tabacum* knockdown (KD) strains; one was a KD for an enzyme of the CBBC that consumes NADPH (glyceraldehyde 3-phosphate dehydrogenase, GAPDH) and a second was a KD for ferredoxin-NADPH oxidoreductase (FNR) which would be limited for its ability to generate NADPH. Both KD strains maintain a similar level of the total NAPDH + NAD^+^ pool, but the GAPDH KD strain has a WT level of NADPH while the pool of pyridine nucleotides in the FNR KD strain is more oxidized (relatively higher levels of NADP^+^). Both KD strains were severely limited for access to oxidized electron acceptors downstream of PSI and exhibited similarly low rates of CO_2_ fixation. While the GAPDH KD strain maintained an oxidized P700 and was therefore protected from photoinhibition, the FNR KD strain was unable to protect PSI from photooxidative damage (Hald et al., 2008). It was hypothesized that redox poise, governed to a large extent by the NADPH/NADP^+^ ratio, regulates electron transport; an elevated NADPH/NADP^+^ ratio would downregulate Cyt *b*_*6*_*f* activity, which would lower the rate of PET and protect PSI from photodamage by favoring oxidized P700 (which is the case in the GAPDH KD) (Hald et al., 2008). FNR, however, binds to Cyt *b*_*6*_*f* and probably forms a complex that facilitates CEF around PSI (Iwai et al., 2010). It has been argued (Joliot and Johnson, 2011) that the H^+^ gradient across the thylakoid membranes, mediated by elevated rates of CEF, is a critical component for eliciting downregulation of Cyt *b*_*6*_*f* activity and preventing PSI photoinhibition. CEF, which competes with LEF and recycles electrons making them unavailable for use by CBBC, would promote the formation of a large H^+^ gradient across the thylakoid membranes (Joliot and Johnson, 2011; Yamamoto and Shikanai, 2019). This large H^+^ gradient could promote the dissipation of excess excitation energy absorbed by PSII as heat (qE component of NPQ), which in turn would diminish the rate of LEF and the generation of ROS. However, some studies suggest that qE may not be an essential component in the protection of PSI from photoinhibition (Joliot and Johnson, 2011; Tikkanen et al., 2015) and Johnson and Berry suggest that the function of qE is to balance the activity of the reaction centers under sub-saturating light conditions for the most effective use of the absorbed light energy in C metabolism (Johnson and Berry, 2021). Additionally, Shimakawa and co-workers (Shimakawa et al., 2018) suggested that the formation of a H^+^ gradient is not sufficient to elicit oxidation (thus protection) of cyanobacterial PSI when CO_2_ levels are low. Finally, this restriction cannot be immediately reversed by uncouplers which would relax the PMF across the thylakoid membranes. If the initial switch is triggered by an elevated PMF and/or redox conditions, then enzymatic processes in the light, whether associated with the ‘opening’ of electron valves downstream of PET and/or the reactivation of the Cyt *b*_*6*_*f* complex through protein modification, may be required (and require some time). Together, these studies suggest the importance of C metabolism/redox poise and potentially the H^+^ gradient across thylakoid membranes in controlling photosynthetic electron flow and Cyt *b*_*6*_*f* complex activity. However, as recently stated (Johnson and Berry, 2021), there is still considerable uncertainty regarding the mechanism(s) of photosynthetic control and the reversible restriction of electron flow across the Cyt *b*_*6*_*f* complex, which may include more than one mechanism for regulating Cyt *b*_*6*_*f* activity.

Similar to a previous study (Shimakawa et al., 2018), our work suggests that the H^+^ gradient is not necessary to sustain restricted electron flow to Cyt *f*. The restricted electron flow to Cyt *f* following dark acclimation of the N-deprived *sta6* mutant was not reversed upon the combined addition of nigericin and valinomycin (**Figure S10**), although as discussed, it was mostly relieved when the mutant was subjected to a short exposure to sub-saturating light.

Despite the restriction in electron transport at the Cyt *b*_*6*_*f* complex, O_2_ production in dark acclimated, N-deprived *sta6* cells was not completely arrested. Real-time measurements of O_2_ exchange demonstrated a maximal O_2_ production rate of 20-25 µmol O_2_ mg Chl^−1^ h^−1^ (this O_2_ was consumed at approximately the same rate as it was generated; **Figure 2**), which is 4-5-fold slower than the rate of O_2_ production by N-deprived C6 cells (∼120 µmol O_2_ mg Chl^−1^ h^−1^; **Figure 6D**). Based on the use of the inhibitor PG, the O_2_ production and uptake (and electron transfer) in N-deprived *sta6* cells could be almost completely attributed to the function of a PTOX dependent H_2_O-to-H_2_O cycle (**Figure 6B** and **C**). Importantly, the rates of O_2_ production and uptake measured when PET was blocked were 5-6 e^−^ PSI^−1^ s^−1^ (conversions between O_2_ production and electron transport rates were performed according to the ratio of 1 e^−^ PSI^−1^ s^−^ ≈ 4 µmol O_2_ mg Chl^−1^ h^−1^, as calculated by Johnson and Alric (Johnson and Alric, 2012)), which correlates with the maximal rate of the PTOX reaction measured in Chlamydomonas (Houille-Vernes et al., 2011), further supporting our proposal that PTOX is the main electron outlet in dark-acclimated, N-deprived *sta6* cells. Therefore, we suggest that the photochemical quenching process mediated by PTOX is an important component of photosynthetic control and may impact PET. The photochemical quenching activity of PTOX may not only contribute to protection of PSII but may also generate a sufficient H^+^ gradient (through the H_2_O-to-H_2_O cycle, which releases 2 H^+^ on the lumenal side of the thylakoid membranes while taking up 2 H^+^ on the stromal side) to enable ATP synthesis and partially fuel the turnover and repair of damaged PSII.

As in previous studies, the work described in this manuscript suggests that photosynthetic control includes the use of an ‘activity’ gauge associated with the conductivity of electrons through the Cyt *b*_*6*_*f* complex or as the Cyt *b*_*6*_*f* was previously described to be functioning like a transistor in an electrical circuit (Johnson and Berry, 2021). Although their work described Cyt *b*_*6*_*f* in controlling steady-state photosynthesis in vascular plants, these results here presented showcase similarities in responses across different organisms and indicate deeply conserved features of this control.

Our results indicate that this control can be activated in the dark in N-deprived *sta6* cells, which generates highly reducing conditions in the chloroplast (and potentially a delta pH across the thylakoid membranes) and be relieved by exposure light (**Figure 3**) through an unidentified electron outlet that supports O_2_ evolution. This restriction helps coordinate the production of low potential electrons with the C assimilatory capacity of the cells and would be especially important for protection of PSI as C fixation saturates. Furthermore, its occurrence in the dark, which would allow N-deprived cells unable to perform starch synthesis to limit PSI damage when the cells become exposed to the light. However, preventing electron flow beyond Cyt *b*_*6*_*f* would increase the reduction state of the PQ pool and possibly cause PSII photodamage. Photosynthetic organisms have evolved mechanisms to rapidly repair PSII and can therefore sustain the ‘cost’ of PSII photodamage while maintaining the benefit of PSI photoprotection. PTOX may also limit PSII damage when PET is restricted; it would reduce PSII photoinhibition by oxidizing a proportion of the PQ pool while also enabling the formation of a H^+^ gradient that may facilitate NPQ and PSII repair. When the cells are once again able to transport photosynthetic electrons (through alternative electron outlets generated when they are kept in the light or when environmental limitations are altered), the restriction is relieved, and electron transport across the Cyt *b*_*6*_*f* complex can proceed.

## METHODS

### Strains

The *sta6* mutant (CC-4348; *sta6*^−^; *rbo1*^−^) (Zabawinski et al., 2001) and the genetically complemented *sta6*::*STA6* strain, designated C6 (CC-4567; *rbo1*^*-*^) (Li et al., 2010) were purchased from the Chlamydomonas Resource Center and maintained under low light on solid Tris-Acetate-Phosphate (TAP) medium. The parental strain for *sta6* and C6 requires arginine for growth and therefore could not be used with our current experimental design.

### Cell growth

Cells were inoculated from solid TAP into liquid TAP medium and grown in flasks at ∼40 μmol photons m^−2^ s^−1^ (aerated by shaking at ∼150 rpm) to mid-logarithmic phase (∼2 × 10^6^ cells mL^−1^). The volume of cell culture containing 240 μg of Chl was pelleted by centrifugation (1,600 x g, 2 min, 21°C), washed twice with fresh photoautotrophic MOPS-Tris medium (20 mM MOPS titrated with TRIS base powder to pH 7.5, and containing 0.2 g L^−1^ K_2_HPO_4_, 0.11g L^−1^ KH_2_PO_4,_ 50 mg L^−1^ CaCl_2_, 100 mg L^−1^ MgSO_4_, 37.5 mg L ^−1^ NH_4_Cl, and 1X Hutner trace elements, modified from (Geraghty et al., 1990)) and re-suspended in the same medium to a final Chl concentration of 4 µg mL^−1^ (∼60 mL of medium). These cultures were bubbled with 2% CO_2_ in glass tubes and grown at 100 μmol photons m^−2^ s^−1^ to a final concentration of ∼10 μg Chl mL^−1^. For N deprivation, photoautotrophically-acclimated cells containing 350-450 μg Chl were pelleted as described above, washed twice with fresh MOPS-TRIS medium devoid of NH_4_Cl, and re-suspended to a final concentration of 5-8 μg Chl mL^−1^ (∼60 mL of medium). These cultures were bubbled with 2% CO_2_ in glass tubes and illuminated with at 100 μmol photons m^−2^ s^−1^ (white light) for 24 h.

### Chl concentration

Chl concentrations were determined from methanol extracted pigments according to (Porra et al., 1989). Briefly, cells were pelleted by centrifugation (13,000 x *g*, 2 min, 21°C) and resuspended in 100% methanol. Cell debris from methanol-extracted samples was pelleted (13,000 x *g*, 5 min, 21°C) and the supernatant containing the pigments was transferred to plastic cuvettes. Total Chl (in µg mL^−1^) was calculated according to the equation: Chl = 22.12 x *A*652 + 2.71 x *A*665, where *A*652 and *A*665 are absorbances at 652 and 665 nm, respectively.

### O_2_ gas exchange

To measure the rates of dark respiration following exposure to different light intensities, the light curves were determined using a Pt-Ag/AgCl polarographic electrode system (ALGI, Littleton, CO, USA) equipped with a temperature-controlled, 1 mL glass water-jacketed reaction chamber, and two YSI 5331A electrodes (Yellow Springs Instruments, Yellow Springs, OH, USA) polarized at -0.8 V. The assay contained 1.5 mL of actively growing cells which were purged with 1% CO_2_/99% He and supplemented with 12 µL of 0.5 M potassium bicarbonate in 5 mM TRIS (pH 7.5). After injection into the reaction chamber, the samples were exposed to stepped light intensities (Luxeon III Star, Lumileds, San Jose, CA, USA) with intensity increases at 3 min intervals followed by 3 min of darkness. The intensities used were 40, 80, 160, 320, 640, 1280, and 1600 µmol photons m^−2^ s^−1^.

### Chl fluorescence analysis

Chl fluorescence was measured using a DUAL-PAM-100 fluorometer (Walz, Germany). For monitoring Chl fluorescence at different light intensities (i.e., a light curve), the cells were supplemented with 2 mM NaHCO_3_ and exposed to increasing light intensities for 1 min at each intensity. The maximal quantum efficiency (Fv/Fm) and actual quantum efficiency of PSII (ΦPSII) as well as photochemical quenching, qL (and 1-qL (Kramer et al., 2004)), which estimate the redox state of the PQ pool with the assumption that Q_A_ and Q_B_ are in equilibrium, were directly calculated from the light curves. For dark acclimation, the cells were vigorously shaken in the dark for 30 min before measuring fluorescence (see **Figure 1C**).

### O_2_ production and light-driven O_2_ photo-reduction

In vivo ^16^O_2_ photoproduction rates and ^18^O_2_ uptake rates were determined using a custom built membrane inlet mass spectrometer (MIMS) with a Pfeiffer quadrupole PrismaPlus QME-220 (Pfeiffer Vacuum, Germany) coupled with a Pt-Ag/AgCl polarographic electrode system (ALGI, Littleton, CO, USA) equipped with a temperature-controlled, water-jacketed 1 mL glass cell, a YSI 5331A electrode (Yellow Springs Instruments, Yellow Springs, OH, USA) polarized at -0.8 V, and an atmospheric barometric pressure sensor (Infineon Technologies Americas Corp, El Segundo, CA, USA). For performing MIMS assays, 1.5 mL of actively growing cells were purged with 1% CO_2_/99% He that was spiked with 0.5 mL 99% ^18^O_2_ and supplemented with 4 mM potassium bicarbonate suspended in fresh medium. After injection of the sample into the reaction chamber, a light (Luxeon III Star, Lumileds, San Jose, CA, USA) regime was applied as follows: 5 min dark, 6 min light (300 µmol photons m^−2^ s^−1^) and then 3 min of dark. The electrode and MIMS were calibrated before each measurement using atmospherically equilibrated medium and 1% CO_2_/99% He purged medium that was spiked with 0.5 mL 99% ^18^O_2_. The light intensity from the LED was calibrated using a Walz US-SQS/L spherical micro quantum sensor (Walz, Germany). Gross O_2_ production and net uptake were calculated according to the following equations (Beckmann et al., 2009)

ΔGross O_2_ uptake = ^18^O_2_ x [1+ (^16^O_2_)/(^18^O_2_)]

ΔGross O_2_ production = ^16^O_2 - ^18^O2_ x [(^16^O_2_)/(^18^O_2_)]

ΔNet O_2_ production = ΔGross O_2_ production – ΔGross O_2_ uptake

To inhibit alternative oxidases, samples were supplemented with 2 mM n-propyl gallate (Sigma) suspended in ethanol. The inhibitor was added just prior to injecting the sample into the glass-reaction vessel.

### Cyt *f* oxidation/reduction kinetics

Laser flash-mediated absorbance changes associated with Cyt *f* oxidation/reduction were monitored using a JTS10 spectrophotometer (Biologic, France) at 554 nm corrected against a base line drawn between the absorption at 546 and 573 nm (de Vitry et al., 2004). N-deprived cells were pelleted by centrifugation (1,600 x g, 2 min, 21°C) and re-suspended to a concentration of 40 µg Chl mL^−1^ in fresh medium supplemented with 10% ficoll (Sigma) and 2 mM NaHCO_3_ (Sigma). Cells were dark acclimated for 30 min before the measurements. For light acclimation, dark-acclimated cells were exposed to 100 µmol photons m^−2^ s^−1^ for the time periods given in the main text. After the acclimation period, the cells were placed in a cuvette in the dark and then oxidation of Cyt *f* was induced by a single turnover laser flash. Absorbance data were normalized to the “a” phase values measured by a laser flash induced electrochromic shift at 520 nm. The half times (T_1/2_) were calculated by fitting the re-reduction phase of the Cyt *f* kinetics curve to a first order decay equation using Origin 2019.

### Oxidation/reduction of P700

Absorbance changes associated with P700 oxidation/reduction were monitored using a JTS10 spectrophotometer (Biologic, France) at 705 nm and corrected using the absorbance at 740 nm (i.e. P700 oxidation/reduction = *ΔI/I* 705 nm - *ΔI/I* 740 nm). Cells in logarithmic growth were pelleted by centrifugation (1,600 x g, 2 min, 21°C) and resuspended to a concentration of 25-30 µg Chl mL^−1^ in fresh medium. P700 oxidation/reduction was monitored over 10 s while the cells were exposed to 60 µmol photons m^−2^ s^−1^, which was followed by a saturating light pulse and a short dark period (as indicated in **Figure 5A**). For dark acclimation, cells were shaken vigorously in darkness for 30 min. For light acclimation, dark-acclimated cells were exposed to 100 µmol photons m^−2^ s^−1^ for 1 h.

### Data analysis

Data were analyzed (statistics, data filtering, and curve fitting) using Origin 2018 Pro and 2019 Pro.

## ACKNOWLEDGMENTS

The authors wish to acknowledge DOE grant DE-SC0008806 and DE-SC0019341, which were awarded to ARG and MCP and which supported most of the work presented in this manuscript. The authors also wish to acknowledge additional support from the Carnegie Institution for Science, which provided SS with part of his salary, and to the Colorado School of Mines for supplying funds to Edward Dempsey for construction of the MIMS system. We had very valuable discussions of our work with Francis-André Wollman, Wojciech Nawrocki, Jennifer E. Johnson, Joe Berry and Xenie Johnson. We would also like to thank other members of the Grossman and Posewitz laboratories for helpful discussions and constructive criticism.

## AUTHOR CONTRIBUTIONS

S.S., M.C.P., and A.R.G. designed research; S.S., D.A.J.K., D.C.T., P.R., and T.M.W. performed the experiments; M.C.P. contributed analytic tools; S.S., D.A.J.K., T.M.W. and A.R.G. analyzed data; and S.S., P.R., M.C.P., and A.R.G. wrote the paper with input from all other coauthors.

## Notes

### Competing Interest Statement

The authors have declared no competing interest.

## REFERENCES

Allahverdiyeva, Y., Mustila, H., Ermakova, M., Bersanini, L., Richaud, P., Ajlani, G., Battchikova, N., Cournac, L., and Aro, E.-M. (2013). Flavodiiron proteins Flv1 and Flv3 enable cyanobacterial growth and photosynthesis under fluctuating light. Proc Natl Acad Sci USA 110: 4111–4116.

Allahverdiyeva, Y., Suorsa, M., Tikkanen, M., and Aro, E.-M. (2015). Photoprotection of photosystems in fluctuating light intensities. J Exp Bot 66: 2427–2436.

Anderson, A., Laohavisit, A., Blaby, I.K., Bombelli, P., Howe, C.J., Merchant, S.S., Davies, J.M., and Smith, A.G. (2016). Exploiting algal NADPH oxidase for biophotovoltaic energy. Plant Biotechnol J 14: 22–28.

Asada, K. (2000). The water–water cycle as alternative photon and electron sinks. Philosophical Transactions of the Royal Society of London. Series B: Biological Sciences 355: 1419–1431.

Bailey, S., Melis, A., Mackey, K.R.M., Cardol, P., Finazzi, G., van Dijken, G., Berg, G.M., Arrigo, K., Shrager, J., and Grossman, A. (2008). Alternative photosynthetic electron flow to oxygen in marine Synechococcus. Biochimica et Biophysica Acta (BBA) - Bioenergetics 1777: 269–276.

Beckmann, K., Messinger, J., Badger, M.R., Wydrzynski, T., and Hillier, W. (2009). On-line mass spectrometry: membrane inlet sampling. Photosynthesis Research 102: 511–522.

Bennoun, P. (1982). Evidence for a respiratory chain in the chloroplast. Proc Natl Acad Sci USA 79: 4352–4356.

Blaby, I.K. et al. (2013). Systems-Level Analysis of Nitrogen Starvation–Induced Modifications of Carbon Metabolism in a Chlamydomonas reinhardtii Starchless Mutant. The Plant Cell 25: 4305–4323.

Buchanan, B.B. (2016). The Path to Thioredoxin and Redox Regulation in Chloroplasts. Annual Review of Plant Biology 67: 1–24.

Chaux, F., Burlacot, A., Mekhalfi, M., Auroy, P., Blangy, S., Richaud, P., and Peltier, G. (2017). Flavodiiron Proteins Promote Fast and Transient O_2_ Photoreduction in Chlamydomonas. Plant Physiology 174: 1825–1836.

Curien, G., Flori, S., Villanova, V., Magneschi, L., Giustini, C., Forti, G., Matringe, M., Petroutsos, D., Kuntz, M., and Finazzi, G. (2016). The Water to Water Cycles in Microalgae. Plant Cell Physiol 57: 1354–1363.

Davey, M.P., Horst, I., Duong, G.-H., Tomsett, E.V., Litvinenko, A.C.P., Howe, C.J., and Smith, A.G. (2014). Triacylglyceride Production and Autophagous Responses in Chlamydomonas reinhardtii Depend on Resource Allocation and Carbon Source. Eukaryotic Cell 13: 392–400.

de Vitry, C. de, Ouyang, Y., Finazzi, G., Wollman, F.-A., and Kallas, T. (2004). The Chloroplast Rieske Iron-Sulfur Protein AT THE CROSSROAD OF ELECTRON TRANSPORT AND SIGNAL TRANSDUCTION. J Biol Chem 279: 44621–44627.

Foyer, C., Furbank, R., Harbinson, J., and Horton, P. (1990). The mechanisms contributing to photosynthetic control of electron transport by carbon assimilation in leaves. Photosynthesis Research 25: 83–100.

Foyer, C. H., Neukermans, J., Queval, G., Noctor, G., and Harbinson, J. (2012). Photosynthetic control of electron transport and the regulation of gene expression. Journal of Experimental Botany, 63: 1637–1661.

Geraghty, A.M., Anderson, J.C., and Spalding, M.H. (1990). A 36 Kilodalton Limiting-CO₂ Induced Polypeptide of Chlamydomonas Is Distinct from the 37 Kilodalton Periplasmic Carbonic Anhydrase. Plant Physiology 93: 116–121.

Gerotto, C., Alboresi, A., Meneghesso, A., Jokel, M., Suorsa, M., Aro, E.-M., and Morosinotto, T. (2016). Flavodiiron proteins act as safety valve for electrons in Physcomitrella patens. PNAS 113: 12322–12327.

Gurrieri, L., Fermani, S., Zaffagnini, M., Sparla, F., and Trost, P. (2021). Calvin–Benson cycle regulation is getting complex. Trends in Plant Science, 26: 898–912.

Hald, S., Nandha, B., Gallois, P., and Johnson, G.N. (2008). Feedback regulation of photosynthetic electron transport by NADP(H) redox poise. Biochimica et Biophysica Acta (BBA) - Bioenergetics 1777: 433–440.

Heinnickel, M., Kim, R.G., Wittkopp, T.M., Yang, W., Walters, K.A., Herbert, S.K., and Grossman, A.R. (2016). Tetratricopeptide repeat protein protects photosystem I from oxidative disruption during assembly. Proc Natl Acad Sci USA 113: 2774–2779.

Houille-Vernes, L., Rappaport, F., Wollman, F.-A., Alric, J., and Johnson, X. (2011). Plastid terminal oxidase 2 (PTOX2) is the major oxidase involved in chlororespiration in Chlamydomonas. Proc Natl Acad Sci USA 108: 20820–20825.

Iwai, M., Takizawa, K., Tokutsu, R., Okamuro, A., Takahashi, Y., and Minagawa, J. (2010). Isolation of the elusive supercomplex that drives cyclic electron flow in photosynthesis. Nature 464: 1210–1213.

Johnson, X. and Alric, J. (2012). Interaction between Starch Breakdown, Acetate Assimilation, and Photosynthetic Cyclic Electron Flow in Chlamydomonas reinhardtii. J Biol Chem 287: 26445–26452.

Johnson, J.E. and Berry, J.A. (2021) The role of Cytochrome b_6_f in the control of steady-state photosynthesis: a conceptual and quantitative model. Photosynth Res 148: 101–136. https://doi.org/10.1007/s11120-021-00840-4.

Joliot, P. and Johnson, G.N. (2011). Regulation of cyclic and linear electron flow in higher plants. Proc Natl Acad Sci USA 108: 13317–13322.

Komenda, J., Sobotka, R., and Nixon, P.J. (2012). Assembling and maintaining the Photosystem II complex in chloroplasts and cyanobacteria. Curr Opin Plant Biol 15: 245–251.

Kono, M., Noguchi, K., and Terashima, I. (2014). Roles of the Cyclic Electron Flow Around PSI (CEF-PSI) and O_2_-Dependent Alternative Pathways in Regulation of the Photosynthetic Electron Flow in Short-Term Fluctuating Light in Arabidopsis thaliana. Plant Cell Physiol 55: 990–1004.

Kozuleva, M., Petrova, A., Milrad, Y., Semenov, A., Ivanov, B., Redding, K. E., and Yacoby, I. (2021). Phylloquinone is the principal Mehler reaction site within photosystem I in high light. Plant Physiology 186:1848–1858.

Kramer, D.M., Johnson, G., Kiirats, O., and Edwards, G.E. (2004). New Fluorescence Parameters for the Determination of QA Redox State and Excitation Energy Fluxes. Photosynthesis Research 79: 209–2018.

Krieger-Liszkay, A. and Feilke, K. (2016). The Dual Role of the Plastid Terminal Oxidase PTOX: Between a Protective and a Pro-oxidant Function. Frontiers in Plant Science https://doi.org/10.3389/fpls.2015.01147.

Li, Y., Han, D., Hu, G., Dauvillee, D., Sommerfeld, M., Ball, S., and Hu, Q. (2010). Chlamydomonas starchless mutant defective in ADP-glucose pyrophosphorylase hyper-accumulates triacylglycerol. Metabolic Engineering 12: 387–391.

Malnoë, A. (2018). Photoinhibition or photoprotection of photosynthesis? Update on the (newly termed) sustained quenching component qH. Environmental and Experimental Botany 154: 123–133.

Mehler, A.H. (1951). Studies on reactions of illuminated chloroplasts: I. Mechanism of the reduction of oxygen and other hill reagents. Archives of Biochemistry and Biophysics 33: 65–77.

Michelet, L., Zaffagnini, M., Morisse, S., Sparla, F., Pérez-Pérez, M.E., Francia, F., Danon, A., Marchand, C.H., Fermani, S., Trost, P., and Lemaire, S.D. (2013). Redox regulation of the Calvin-Benson cycle: something old, something new. Front Plant Sci 4: 470.

Müller, P., Li, X.-P., and Niyogi, K.K. (2001). Non-Photochemical Quenching. A Response to Excess Light Energy. Plant Physiology 125: 1558–1566.

Munekage, Y., Hojo, M., Meurer, J., Endo, T., Tasaka, M., and Shikanai, T. (2002). PGR5 Is Involved in Cyclic Electron Flow around Photosystem I and Is Essential for Photoprotection in Arabidopsis. Cell 110: 361–371.

Munekage, Y.N., Genty, B., and Peltier, G. (2008). Effect of PGR5 Impairment on Photosynthesis and Growth in Arabidopsis thaliana. Plant Cell Physiol 49: 1688–1698.

Nath, K., Jajoo, A., Poudyal, R.S., Timilsina, R., Park, Y.S., Aro, E.-M., Nam, H.G., and Lee, C.-H. (2013). Towards a critical understanding of the photosystem II repair mechanism and its regulation during stress conditions. FEBS Letters 587: 3372–3381.

Nawrocki, W.J., Tourasse, N.J., Taly, A., Rappaport, F., and Wollman, F.-A. (2015). The Plastid Terminal Oxidase: Its Elusive Function Points to Multiple Contributions to Plastid Physiology. Annual Review of Plant Biology 66: 49–74.

Nixon, P.J. (2000). Chlororespiration. Philosophical Transactions of the Royal Society of London. Series B: Biological Sciences 355: 1541–1547.

Peltier, G. and Cournac, L. (2002). Chlororespiration. Annual Review of Plant Biology 53: 523–550.

Porra, R.J., Thompson, W.A., and Kriedemann, P.E. (1989). Determination of accurate extinction coefficients and simultaneous equations for assaying chlorophylls a and b extracted with four different solvents: verification of the concentration of chlorophyll standards by atomic absorption spectroscopy. Biochimica et Biophysica Acta (BBA) - Bioenergetics 975: 384–394.

Roach, T. and Krieger-Liszkay, A.K. (2014). Regulation of Photosynthetic Electron Transport and Photoinhibition. Curr Protein Pept Sci 15: 351–362.

Rochaix, J.-D. (2014). Regulation and Dynamics of the Light-Harvesting System. Annual Review of Plant Biology 65: 287–309.

Sarewicz, M., Bujnowicz, Ł., Bhaduri, S., Singh, S.K., Cramer, W.A., and Osyczka, A. (2017). Metastable radical state, nonreactive with oxygen, is inherent to catalysis by respiratory and photosynthetic cytochromes bc1/b6f. Proc Natl Acad Sci USA 114: 1323–1328.

Saroussi, S., Karns, D.A.J., Thomas, D.C., Bloszies, C., Fiehn, O., Posewitz, M.C., and Grossman, A.R. (2019). Alternative outlets for sustaining photosynthetic electron transport during dark-to-light transitions. Proc Natl Acad Sci USA 116: 11518–11527.

Saroussi, S.I., Wittkopp, T.M., and Grossman, A.R. (2016). The Type II NADPH Dehydrogenase Facilitates Cyclic Electron Flow, Energy-Dependent Quenching, and Chlororespiratory Metabolism during Acclimation of Chlamydomonas reinhardtii to Nitrogen Deprivation1[OPEN]. Plant Physiol 170: 1975–1988.

Schmollinger, S. et al. (2014). Nitrogen-Sparing Mechanisms in Chlamydomonas Affect the Transcriptome, the Proteome, and Photosynthetic Metabolism. The Plant Cell 26: 1410–1435.

Shaku, K., Shimakawa, G., Hashiguchi, M., and Miyake, C. (2016). Reduction-Induced Suppression of Electron Flow (RISE) in the Photosynthetic Electron Transport System of Synechococcus elongatus PCC 7942. Plant Cell Physiol 57: 1443–1453.

Shimakawa, G., Shaku, K., and Miyake, C. (2016). Oxidation of P700 in Photosystem I Is Essential for the Growth of Cyanobacteria. Plant Physiology 172: 1443–1450.

Shimakawa, G., Shaku, K., and Miyake, C. (2018). Reduction-Induced Suppression of Electron Flow (RISE) Is Relieved by Non-ATP-Consuming Electron Flow in Synechococcus elongatus PCC 7942. Front. Microbiol. 9.

Shubin, V.V., Terekhova, I.N., Kirillov, B.A., and Karapetyan, N.V. (2008). Quantum yield of P700+ photodestruction in isolated photosystem I complexes of the cyanobacterium Arthrospira platensis. Photochem. Photobiol. Sci. 7: 956–962.

Siaut, M., Cuiné, S., Cagnon, C., Fessler, B., Nguyen, M., Carrier, P., Beyly, A., Beisson, F., Triantaphylidès, C., Li-Beisson, Y., and Peltier, G. (2011). Oil accumulation in the model green alga Chlamydomonas reinhardtii: characterization, variability between common laboratory strains and relationship with starch reserves. BMC Biotechnology 11: 7.

Smith, R. T., and Gilmour, D. J. (2018). The influence of exogenous organic carbon assimilation and photoperiod on the carbon and lipid metabolism of Chlamydomonas reinhardtii. Algal Research, 31: 122–137.

Sonoike, K. (1996). Degradation of psaB gene product, the reaction center subunit of photosystem I, is caused during photoinhibition of photosystem I: possible involvement of active oxygen species. Plant Science 115: 157–164.

Sonoike, K. (2011). Photoinhibition of photosystem I. Physiologia Plantarum 142: 56–64.

Suorsa, M., Järvi, S., Grieco, M., Nurmi, M., Pietrzykowska, M., Rantala, M., Kangasjärvi, S., Paakkarinen, V., Tikkanen, M., Jansson, S., and Aro, E.-M. (2012). PROTON GRADIENT REGULATION5 Is Essential for Proper Acclimation of Arabidopsis Photosystem I to Naturally and Artificially Fluctuating Light Conditions. The Plant Cell 24: 2934–2948.

Terashima, I., Funayama, S., and Sonoike, K. (1994). The site of photoinhibition in leaves of Cucumis sativus L. at low temperatures is photosystem I, not photosystem II. Planta 193: 300–306.

Terashima, I., Noguchi, K., Itoh-Nemoto, T., Park, Y.-M., Kuhn, A., and Tanaka, K. (1998). The cause of PSI photoinhibition at low temperatures in leaves of Cucumis sativus, a chilling-sensitive plant. Physiologia Plantarum 103: 295–303.

Theis, J. and Schroda, M. (2016). Revisiting the photosystem II repair cycle. Plant Signal Behav 11(9): e1218587.

Tikhonov, A. N. (2014). The cytochrome b6f complex at the crossroad of photosynthetic electron transport pathways. Plant Physiology and Biochemistry 81: 163–183.

Tikhonov, A. N. (2015). Induction events and short-term regulation of electron transport in chloroplasts: an overview. Photosynthesis Research 125: 65–94.

Tikkanen, M., Rantala, S., and Aro, E.-M. (2015). Electron flow from PSII to PSI under high light is controlled by PGR5 but not by PSBS. Front Plant Sci 6.

Weis, E., Ball, J.T., and Berry, J. (1987). Photosynthetic Control of Electron Transport in Leaves of Phaseolus Vulgaris: Evidence for Regulation of Photosystem 2 by the Proton Gradient. In Progress in Photosynthesis Research: Volume 2 Proceedings of the VIIth International Congress on Photosynthesis Providence, Rhode Island, USA, August 10–15, 1986, J. Biggins, ed (Springer Netherlands: Dordrecht), pp. 553–556.

Wirtz, W., Stitt, M., and Heldt, H.W. (1982). Light activation of calvin cycle enzymes as measured in pea leaves. FEBS Letters 142: 223–226.

Xu, C., Moellering, E. R., Fan, J., and Benning, C. (2008). Mutation of a mitochondrial outer membrane protein affects chloroplast lipid biosynthesis. The Plant Journal, 54, 163–175.

Yamamoto, H. and Shikanai, T. (2019). PGR5-Dependent Cyclic Electron Flow Protects Photosystem I under Fluctuating Light at Donor and Acceptor Sides. Plant Physiology 179: 588–600.

Yamori, W., Makino, A., and Shikanai, T. (2016). A physiological role of cyclic electron transport around photosystem I in sustaining photosynthesis under fluctuating light in rice. Sci Rep 6: 20147.

Zabawinski, C., Koornhuyse, N.V.D., D’Hulst, C., Schlichting, R., Giersch, C., Delrue, B., Lacroix, J.-M., Preiss, J., and Ball, S. (2001). Starchless Mutants of Chlamydomonas reinhardtii Lack the Small Subunit of a Heterotetrameric ADP-Glucose Pyrophosphorylase. Journal of Bacteriology 183: 1069–1077.

Zivcak, M., Brestic, M., Kunderlikova, K., Sytar, O., and Allakhverdiev, S.I. (2015). Repetitive light pulse-induced photoinhibition of photosystem I severely affects CO_2_ assimilation and photoprotection in wheat leaves. Photosynthesis Research 126: 449–463.

