## Supplementary Information for "Reversible restriction of electron flow across cytochrome *b_6_f* in dark acclimated cells limited for downstream electron sinks"

*Corresponding authors:

Arthur Grossman

### Material and Methods

**Starch accumulation:** C6 and *sta6* cells were pelleted by centrifugation (1600 x g, 2 min, 21°C) and resuspended in 100% methanol. The extracted sample was centrifuged (maximal speed in the microfuge, 1 min, RT), the supernatant discarded, and the pellet (including the starch granules) was resuspended in 0.5 mL $H_2O$ and autoclaved for 20 min followed by the addition of 1 µl of 0.01 N potassium iodide to 5 µL of the autoclaved pellet. Starch was further qualitatively measured by visual inspection.

**Assessment of P700 re-reduction rate**: Absorbance changes associated with P700 oxidation/reduction were monitored using a JTS10 spectrophotometer (Biologic, France) at 705 nm and corrected according to the absorbance at 740 nm (i.e., P700 oxidation/reduction = $\Delta I/I$ 705 nm - $\Delta I/I$ 740 nm). Cells in logarithmic growth were pelleted by centrifugation (1600 x g, 2 min, at 21°C) and resuspended to a concentration of 25-30 µg $mL^{-1}$ of Chl in fresh medium. P700 oxidation/reduction of dark-acclimated cells (30 min) was monitored over 10 s while the cells were exposed to 156 µmol photons $m^{-2}$ $s^{-1}$ followed by a pulse of a saturating light and then 5 s of darkness. 20 µM DCMU and 1 mM hydroxylamine were included in the assay to inhibit LEF. The re-reduction rate of P700 was calculated according to [1,2].

**Dark respiration:** Dark respiration at different light intensities was measured using a Pt-Ag/AgCl polarographic electrode system (ALGI, Littleton, CO, USA) equipped with a temperature-controlled, 1 mL glass water-jacketed reaction chamber, two YSI 5331A electrodes (Yellow Springs Instruments, Yellow Springs, OH, USA) polarized at -0.8 V, and an atmospheric barometric pressure sensor (Infineon Technologies Americas Corp, El Segundo, CA, USA). Assays were performed using 1.5 mL of actively growing cells that were purged with 1% $CO_2$/99% He supplemented with 12 µL of 0.5 M potassium bicarbonate in 5 mM Tris. After injection into the reaction chamber, the rate of change in $O_2$ levels was measured sequentially at light intensities of 40, 80, 160, 320, 640, 1280, and 1600 µmol photons $m^{-2}$ $s^{-1}$ (PAR) (Luxeon III Star, Lumileds, San Jose, CA, USA); each intensity was maintained for 3 min followed by a 3 min intervening dark period and then the light level was raised to the next higher intensity (step change) until the full range of intensities were tested. Prior to the measurements for each series of light intensities, the electrodes were calibrated with air (~21% $O_2$) and with 1% $CO_2$/99% He purged MOPS medium (0% $O_2$). The intensity of light from the LEDs was calibrated using a Walz US-SQS/L spherical micro quantum sensor (Walz, Germany).

**Protein extraction and immunoblot detection:** Total cell protein was obtained by pelleting cells by centrifugation (1,200xg, 5 min, and 23°C), resuspending the pellets in a protein extraction buffer which contained 5 mM HEPES-KOH, pH 7.5, 100 mM dithiothreitol, 100 mM $Na_2CO_3$, 2% (w/v) SDS, and 12% (w/v) sucrose, and boiling the samples for 1 min. Samples used for SDS-PAGE (Criterion 12% acrylamide gels, Bio-Rad) were normalized to cell number (and similar levels of tubulin), and the resolved polypeptides were transferred to polyvinylidene difluoride membranes using the Trans-Blot Turbo Transfer System (Bio-Rad) according to the manufacturer's directions. For immunoblot analysis, membranes were blocked for 1 h at room temperature in Tris-buffered saline-0.1% (v/v) Tween containing 5% (w/v) milk powder followed by an overnight incubation of the membranes at 4°C with the primary antibodies in Tris-buffered saline-0.1% (v/v) plus Tween20 containing 3% (w/v) milk powder. Primary antibodies against D1, Tubulin, and Cyt *f* were diluted according to the manufacturer's suggestions (Agrisera), while the antibodies against PC (gift from the laboratory of Sabeeha Merchant) were diluted 1:500 in Tris-buffered saline-0.1% (v/v) plus Tween20 containing 3% (w/v) milk powder. Proteins were detected by enhanced chemiluminescence (GE Healthcare).

### Supplementary Figures

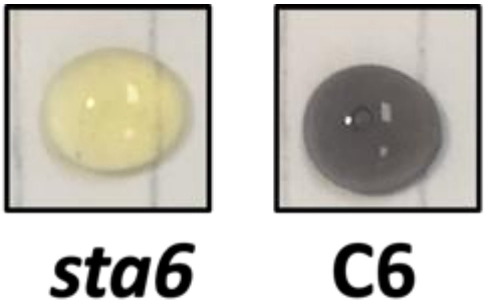


**Figure S1: Qualitative starch assay:** Iodine assay for the presence of starch in both the starchless mutant, *sta6* (yellowish color, absence of starch), and the genetically complemented C6 strain (dark purple color, presence of starch).

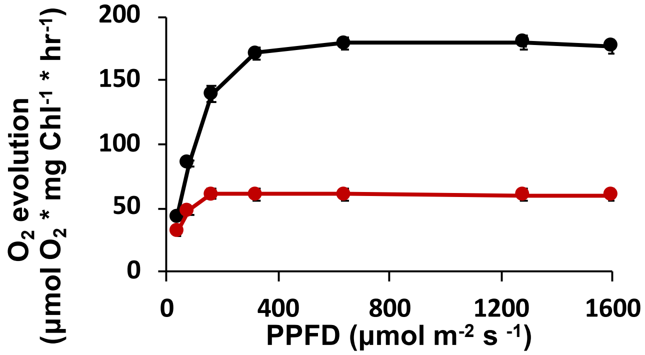


**Figure S2: $O_2$ evolution in the presence of N:** Net $O_2$ evolution of N-replete C6 (black curve) and *sta6* (red curve). N=3 ± SE.

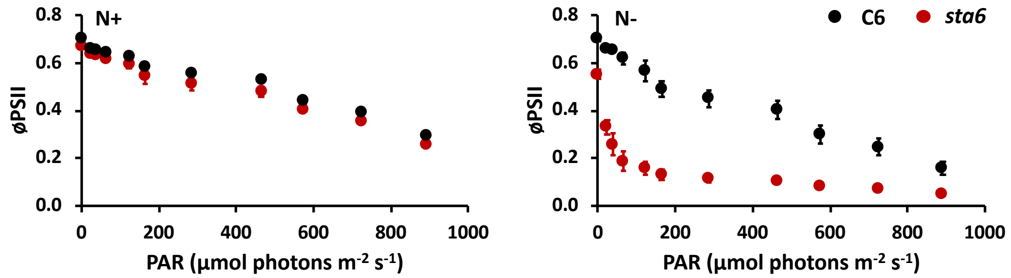


**Figure S3**: **The effective quantum yield (ΦPSII) under N-replete and N-deprivation growth conditions:** Measurements of photosynthetic yield for C6 and *sta6* cultured in the presence (**A**) and absence (**B**) of N. For all panels, data for C6 is presented in black while data for *sta6* is in red. ΦPSII was measured as a function of light while *sta6* (red circles) and C6 (black circles) cells were cultured in the presence (left panel) and absence (right panel) of N in the medium. The first data point represents the maximal quantum yield (Fv/Fm). N=3 ± S.D.

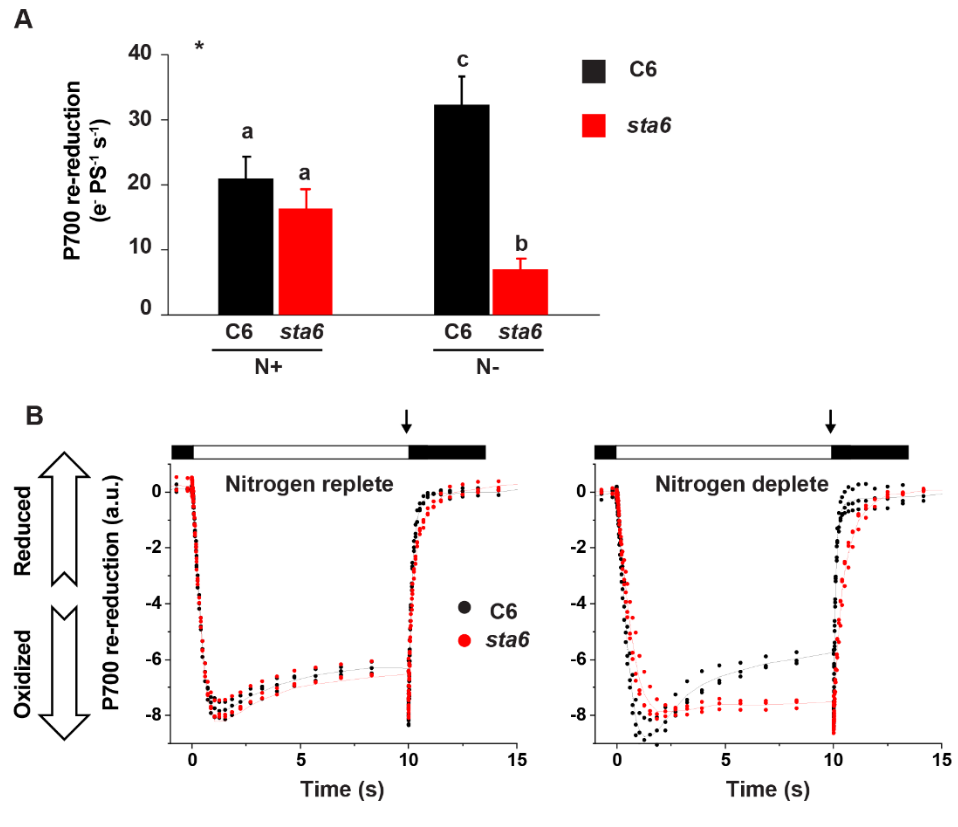


**Figure S4**: **P700 re-reduction rate and oxidation/reduction kinetics following dark acclimation: A.** Measurements were performed in the presence of 20 µM DCMU and 1 mM hydroxylamine to inhibit LEF. Rates were calculated as indicated in the Method section using kinetics similar to those shown in the **B** panels. * One -way ANOVA (*p*-value = 0.01) followed by a Tuky post-hoc test (*p*-value = 0.01). N=4-6 ± S.D. **B.** P700 oxidation/reduction kinetics that were used for determining the rates of re-reduction. Cells were cultured in the presence (left) and absence (right) of N. Saturating pulses were applied immediately before the cells were transitioned from light to dark, as indicated by the black arrow. N=3 ± S.D.

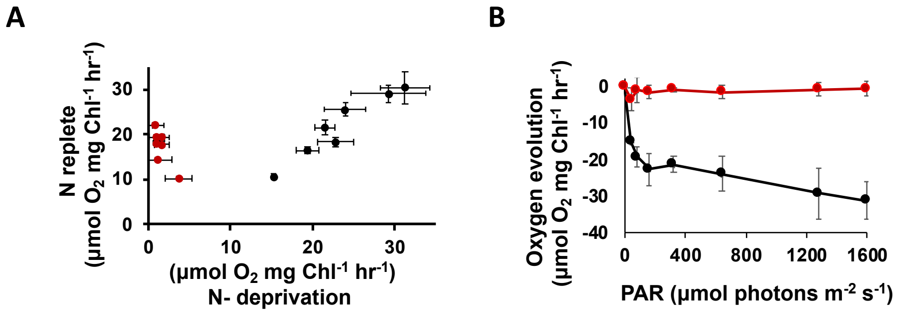


**Figure S5: $O_2$ gas exchange. A.** Correlation between dark respiration rates measured in the presence and absence of N upon exposure of C6 (black) and *sta6* (red) to increasing light intensities; 40, 80, 160, 320, 640, 1280, and 1600 µmol photons $m^{-2}$ $s^{-1}$. **B.** Dark respiration of N-deprived C6 (black) and *sta6* (red) indicated as $O_2$ uptake (higher uptake rates have a more negative value) following exposure to increasing light intensities. N=3 ± S.D.

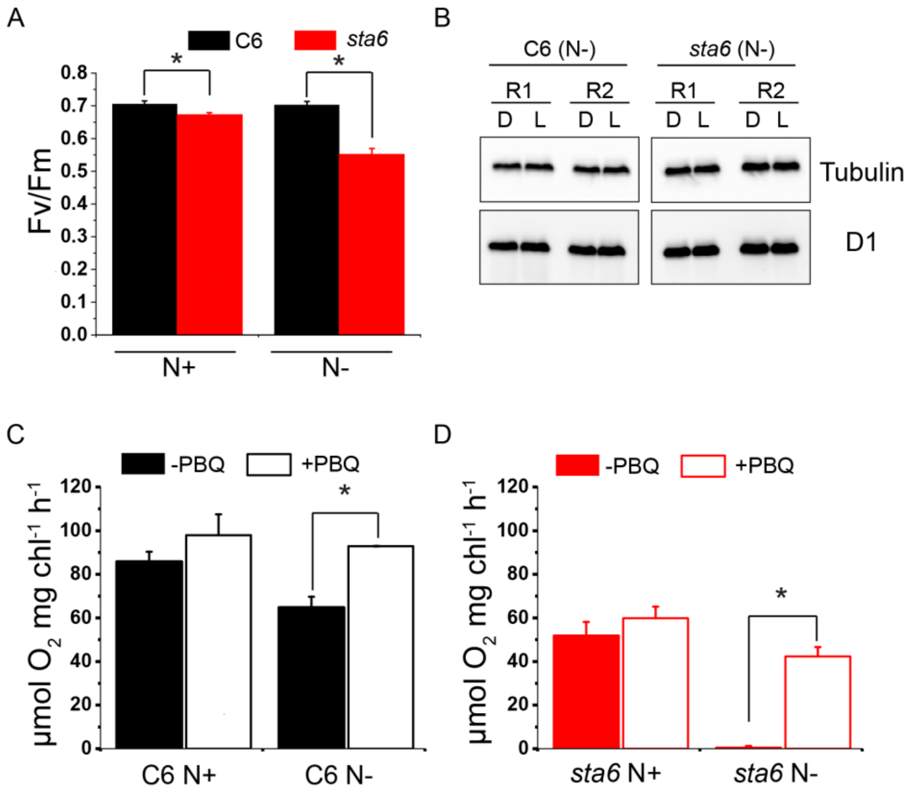


**Figure S6**: **PSII activity and the level of the D1 protein: A**. Maximal quantum yield (Fv/Fm) for N-replete (N +, left pair of bars) and N-deprived (N-, right pair of bars) C6 and *sta6*. * t-test, *p*-value < 0.05 N =3 ± SD. **B.** Immunoblot of D1 protein from light- (L) and dark- (D) acclimated N-deprived (N-) C6 and *sta6* cells. Two biological replicates (R) from each strain are shown. Blots were taken from the same gel. **C** and **D**: Net $O_2$ production for C6 (**C**) and *sta6* (**D**) in the presence and absence of 80 µM *p*-benzoquinone (PBQ). * t-test, *p*-value=0.05 N =3 ± SD.

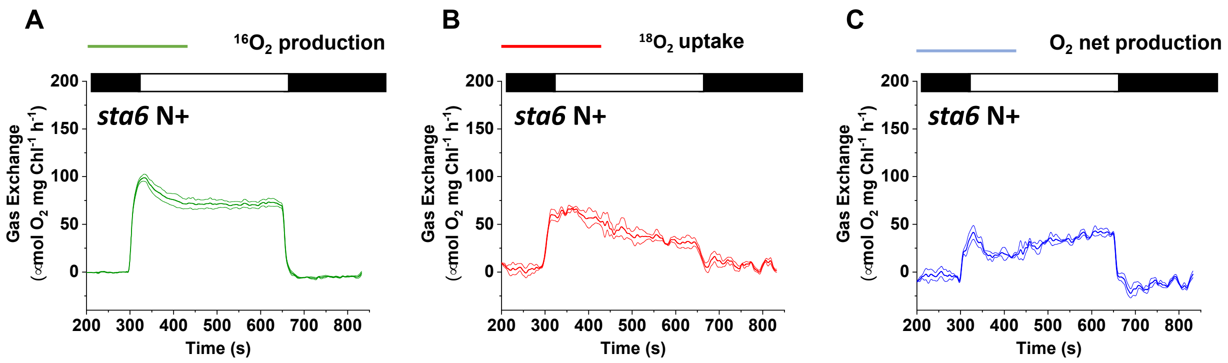


**Figure S7: Real-time $O_2$ exchange for N-replete *sta6* using MIMS:** Light-induced gross $O_2$ production ($^{16}O_2$ evolution, green curve) **A**, and uptake ($^{18}O_2$ consumption, red curve) **B**, for N-replete *sta6* following a dark (30 min) to light (300 µmol photons $m^{-2}$ $s^{-1}$) transition; gas exchange was monitored for 6 min in the light using MIMS. Prior to turning on the light, the cells were maintained for 5 min in the dark (in the MIMS chamber) and then for 3 min in the dark following illumination. Net $O_2$ production (blue curve), **C**, was calculated according to the formula given in Methods section of the main text. Each trace represents an average of at least 3 biological replicates ± SE.

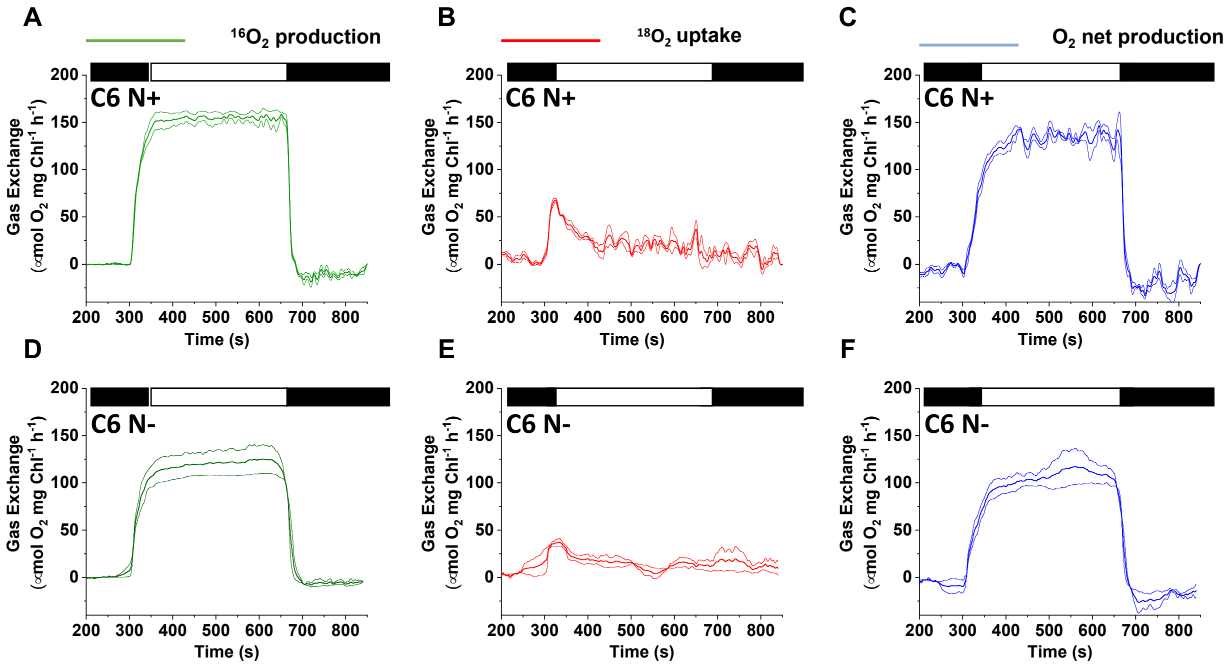


**Figure S8**: **Real-time $O_2$ exchange for C6 cells using MIMS:** Light-induced gross $O_2$ production ($^{16}O_2$ evolution, green curve), and uptake ($^{18}O_2$ consumption, red curve), for N-replete (**A** and **B**, respectively) and N-deplete (**D** and **E**, respectively) C6 cells following a dark (30 min) to light (300 µmol photons $m^{-2}$ $s^{-1}$) transition; gas exchange was monitored for 6 min in the light using MIMS. Prior to turning on the light, the cells were maintained for 5 min in the dark (in the MIMS chamber) and then for 3 min in the dark following illumination. Net $O_2$ production (blue curve), was calculated for N-replete (**C**) and N-deplete (**F**) C6 cells according to the formula given in **Materials** section of the main text. Each trace represents an average of at least 3 biological replicates ± SE.

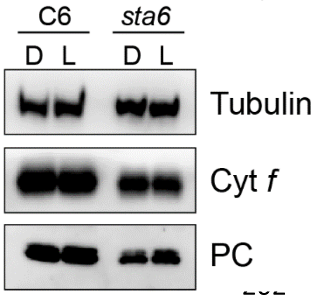


**Figure S9:** Immunoblot analysis of Cyt *f* subunit of the Cyt $b_6f$ complex and PC. Total protein extracts derived from the same number of cells were loaded onto a 12% polyacrylamide gel and resolved by SDS-PAGE. A monospecific antibody was used to detect Cyt *f*, PC, and β-tubulin; tubulin was used as a loading control.

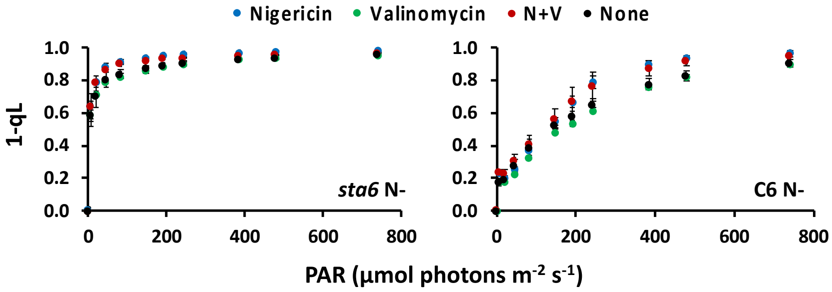


**Figure S10**: **Impact of ΔpH and ΔΨ on 1-qL:** 1-qL was calculated for N-deprived (for 24 h) *sta6* (left panel) and C6 cells (right panel) as shown in **Figure 3** in the main text. Analyses were performed in the absence of inhibitors (black circles), presence of 10 μM nigericin (blue circles), presence of 10 μM valinomycin (green circles), or in the presence of both nigericin and valinomycin (N+V, red circles). n=3 ± S.D.

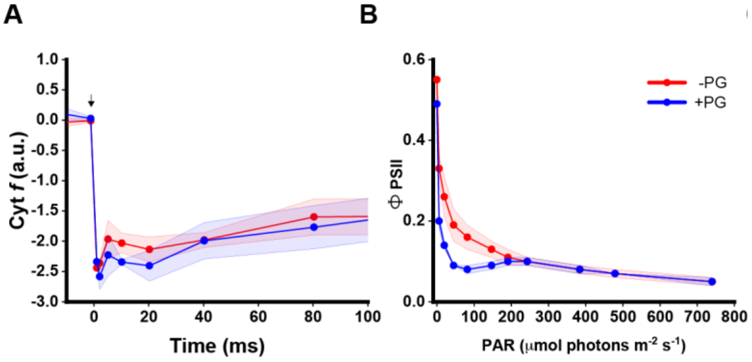


**Figure S11: The effect of the PTOX inhibitor, PG, on photosynthetic electron transport reactions in the N-deprived *sta6* mutant:** Following dark acclimation (30 min), the samples were supplemented with 2 mM PG and incubated in the dark for an additional 5 min. **A.** Oxidation/reduction kinetics of Cyt *f*. Experimental design was as in **Figures 4A** and **B** in the main text. The black arrow indicates administration of a saturating laser pulse. **B**. PSII efficiency.

**Reference:**


1. Clowez, S., Godaux, D., Cardol, P., Wollman, F.-A. & Rappaport, F. The Involvement of Hydrogen-producing and ATP-dependent NADPH-consuming Pathways in Setting the Redox Poise in the Chloroplast of *Chlamydomonas reinhardtii* in Anoxia. *J. Biol. Chem.* **290**, 8666–8676 (2015).
2. Takahashi, H., Clowez, S., Wollman, F.-A., Vallon, O. & Rappaport, F. Cyclic electron flow is redox-controlled but independent of state transition. *Nat. Commun.* **4**, 1–8 (2013).
